# Ontogenetic sequence of differential gene expression in predator-induced *Daphnia pulex*

**DOI:** 10.1101/2025.07.16.665143

**Authors:** Andrey Rozenberg, Linda C. Weiss, Tatjana Schwarz, Nancy Kühne, Uwe John, Ralph Tollrian

## Abstract

As a reaction to the presence of predators, many *Daphnia* species develop morphological defenses. These defenses are phenotypically plastic traits as their production is switched off in the absence of the predator. Previous studies identified some neurohumoral factors and candidate genes that are involved in the development of the defenses. However, there is still significant uncertainty regarding the involvement of certain genes and factors, and minimal information is available about the timing of gene expression changes throughout this development. In the current study, we thus performed a candidate-independent gene expression analysis of defense development over juvenile developmental stages in *D. pulex*. We analyzed transcriptome responses of the microcrustaceans to the natural mix of predator-emitted compounds and also to the pure kairomone which had been identified recently and which we synthesized. With the data obtained we show that the main factor correlating with the global gene expression patterns in *Daphnia* is molt cycle. Influence of kairomone treatment on the level of global gene expression is evident only in certain stages. The stages generally show unique patterns of kairomone-induced gene expression. However, expression profiles are highly similar for the naturally released and the chemically synthesized kairomone in each one of the stages. A number of genes with regulatory, structural and detoxification roles are differentially expressed as a reaction to the kairomone treatment. The most consistent response was found in the expression levels of *ilp-3* coding for an insulin-like peptide. Gene knockdown experiments suggest that this hormone plays a role in the production of the defense. Many of the genes responding to the kairomone treatment have no predicted function stressing the need to investigate gene functions in *Daphnia*.

## Introduction

*Daphnia* micro-crustaceans adapt their life-cycle, behavior and/or morphology when exposed to predation (1). Most of these changes act as protection against predator attacks, but their production comes at a cost, which is saved in the absence of the predator (2,3). *Daphnia* defensive mechanisms are very diverse and often involve easily detectable morphological alterations in the form of elongated spines, helmets, crests, as well as small cuticular neckteeth or peculiar “crown of thorns” (1,4). Some of these morphological structures emerged several times in the evolution of the genus. Thus, the neckteeth developed under predation by the phantom midge larvae *Chaoborus* are characteristic to several not directly related lineages of *Daphnia*, including the *D. pulex* group (1,5–7).

Information about the predator’s presence is mediated mainly by signalling cues unintentionally produced and released by the predator, referred to as kairomones (1). The chemical nature of most kairomones capable of inducing physiological reactions in *Daphnia* remains unknown, but attempts at isolation and characterization of kairomones with physiological effects on *Daphnia* have been undertaken since the 1990s (8–11). Among the few kairomones whose precise chemical structures have been elucidated and functionally validated in physiological experiments, one of the best-characterized is the fish kairomone 5α-cyprinol sulfate. This bile salt was identified as the active compound inducing diel vertical migration in *Daphnia* (12), and represents a rare example of a structurally defined predator cue. Similarly, the kairomones emitted by *Chaoborus* have been chemically identified and experimentally validated (13). The *Chaoborus* kairomone consists of a family of long-chained (≥C14) fatty acids coupled to the α-amine of L-glutamine. We previously identified six structurally similar compounds within this mixture that are biologically active and induce neckteeth formation in *Daphnia pulex* (13). Among them, N-oleoyl L-glutamine was the most abundant, although all identified components showed comparable potency.

Notwithstanding the diversity of the defensive structures, predators and their kairomones, all of the *Daphnia* morphological defenses are associated with a specific population of large polyploid cells in the corresponding head regions (14–16). Moreover, the same set of neurohumorally active compounds: dopamine and acetylcholine, is involved in the formation of the structurally dissimilar neckteeth and crests in *D. pulex* and *D. longicephala*, respectively (15). At least in *D. pulex*, juvenile hormone (17,18) is seemingly also involved in the regulation of the neckteeth production, although a recent study failed to reproduce results of some of the earlier qPCR experiments (19). Based on gene expression data, the life history shifts accompanying the neckteeth production were connected to ecdysone (18,20), although the role of ecdysone was also questioned by Christjani et al. (19). Neckteeth development and the overall thickening of the cuticle were shown to be correlated with up-regulation of several cuticle-associated proteins (19,21–23), even though temporal and spatial patterns of their expression remain to be explored. Little is known about the early steps of kairomone perception: glutamate receptors were shown to be up-regulated in the embryos shortly after kairomone exposure (23,24) and their antagonists were shown to inhibit formation of the neckteeth (24), but the kairomone receptor itself remains unidentified.

Given the conflicting evidence of the role of some of the neurohumoral factors in the production of the defenses even in the best studied system *D. pulex*–*Chaoborus* (19), as well as the fact that every next study finds new regulators and new target genes i.e. (22,24), one might conclude that the understanding of defense development is far from complete. Moreover, temporal patterns of gene expression might offer a solution to the conflicting reports. These considerations motivated us to perform a replicated time-series study of gene expression using a candidate-independent approach. We chose to focus on four distinct time points covering the periods from kairomone perception to the presumptive decline of its effect (Fig. 1a). As other compounds co-produced by the predator along the kairomone, as well as alarm cues from the prey (25) might be physiologically active and consequently influence genetic responses, we decided to perform two sets of treatment experiments: the kairomone as part of the natural mix of predator-associated compounds and the chemically synthesized pure kairomone. To increase statistical power of the differential gene expression analysis, each combination of treatment/stage was performed in six biological replicates, with libraries prepared for individual clutches. The focus on single clutches allowed us to achieve a precise synchronization between individuals in the same sample, which, as we demonstrate, is crucial to accounting for life-history changes. Gene expression assessment was carried out following our protocol developed for Tag-Seq (26).

**Figure 1.**
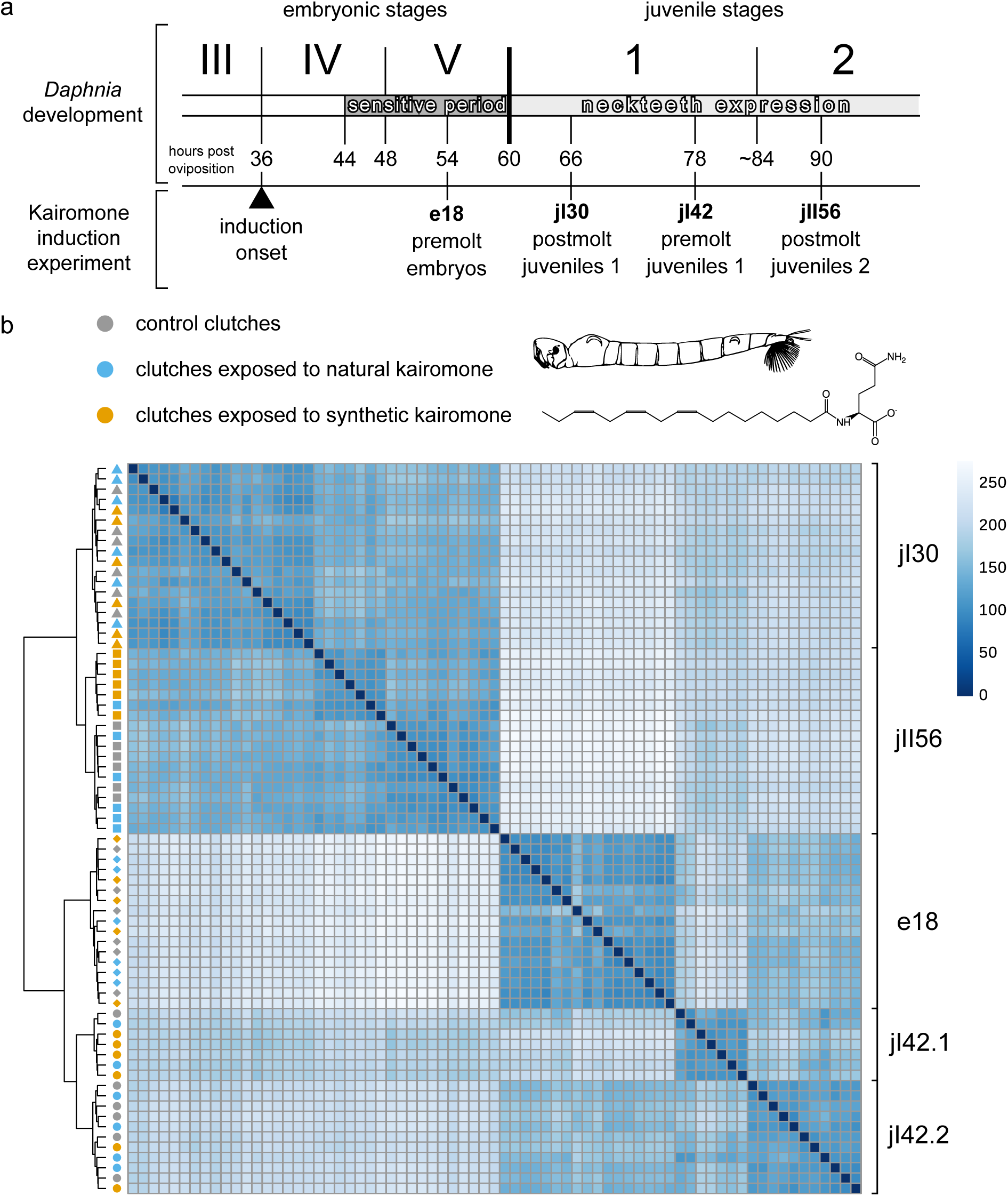
An overview of the kairomone-induction experiment used to generate Tag-Seq libraries. **a**, Experimental setup utilized in the current study in the context of early development of *D. pulex*. Morphological stages are classified according to (27) and are based on easily-recognizable landmarks, but do not directly correspond to intervals between embryonic molts (28,99) shown above as dashed arrows. Solid arrows denote post-embryonic exuviations, the first of which occurs within 20–30 min after neonates leave their mother’s brood pouch (100,101). The “kairomone timing” bar shows kairomone-sensitive periods (28,102) (dark gray) and the period of neckteeth expression (light gray). Experiments performed in this study were initiated with the appearance of eyes in embryos, 8 hours before the onset of the sensitive period. Four time points were sampled after 18 hours with an interval of 12 hours. A single sample corresponded to a single clutch of siblings. **b**, Heatmap and hierarchical clustering of the *Daphnia* Tag-Seq libraries according to overall expression patterns based on data mapped to the R9 assembly and rlog-normalized with DESeq2. Sample names denote stage (“e” for embryo, “jI” for juvenile 1 and “jII” for juvenile 2), time since induction onset in hours, and experimental condition. The outlier sample e18fs1 is omitted from the plot.

## Material and methods

### Material and experimental design

The clone R9 used for the experiments belongs to the species in the literature referred to as North American or Pan-Arctic *Daphnia “pulex”* and was originally isolated from the Ontario province, Canada. The general scheme of the experiment and its relation to *D. pulex* development are shown in Figure 1. Animals were propagated strictly parthenogenetically in 10 L buckets with aged charcoal-filtered tap water under constant light and fed with *Chlorella vulgaris* micro-algae *ad libitum* (<1.5 gC/L). For seven generations before the onset of the experiment, every next generation was initiated from juveniles of the third or later clutch to decrease inter-individual variation. After the females of the eighth generation released their third-fourth clutch, their reproductive cycle was monitored on an individual basis. 72 females in total were selected when embryos in their brood pouches reached the early fourth stage (27), which is a morphologically easily recognizable time point eight hours before the onset of the kairomone-sensitive period (28) (Fig. 1). Each female was placed individually in a 50 ml glass vial with one of the three following media:

1. Control: same water as used for the culture, void of predator cues.
2. Natural kairomone: 1:5 dilution of phantom midge larvae-exposed water in the water used for the culture. The stock solution was prepared by placing 50 *Chaoborus* larvae fed on 500 juvenile R9 *Daphnia* in a one-liter jar for 24 hrs. The stock was prepared in two batches, filtered using a GF/C-grade filter, mixed and frozen as one-time aliquots at –20 °C until usage.
3. Synthetic kairomone (13): 1:25000 dilution of a stock solution of N-linolenoyl-L-glutamine. The stock was prepared by dissolving the chemically synthesized compound in a 1:10 dioxane:ethanol mixture to a concentration of 12.2 mM.

As the synthetic kairomone medium contained minute amounts of dioxane and ethanol, equal volume of the same solvent mixture was also added to the two other media. For nourishment, *C. vulgaris* to the final concentration of 1.5 mg carbon/l was added in each vial (stock concentrations were calculated by calibrated photometric measurements). All media were mixed well before placing the animals.

24 hrs after the onset of the experiment females released the neonates (see Fig. 1), which were placed in new vials one clutch per vial, with fresh media having the same amounts of the components. The juveniles of the first instar molted to the second instar roughly after 24 hrs post-release.

The experimental animals were sampled at four time points (see Fig. 1): pre-release embryos 18 hrs after induction onset (e18), juveniles of the 1^st^ instar after 30 hrs (jI30), juveniles of the 1^st^ instar after 42 hrs (jI42) and juveniles of the 2^nd^ instar after 56 hrs (jII56). Each of the treatment-stage combinations was replicated six times, with individual clutches assigned as random as possible.

Before fixation, the juveniles were controlled for the absence or presence of the neckteeth in the control and induction media, respectively. Each clutch was fixed separately in the RLT buffer from the RNeasy Micro Kit (Qiagen). Mean (±SD) number of progeny per clutch amounted to 21.1±4.5, but no more than 18 individuals were sampled to reduce the variation between the clutches (resulting in 17.2±1.9 individuals per clutch sampled). Upon fixation, the tubes were stored at 4 °C overnight and frozen away at –80 °C afterwards.

### Tag-Seq library preparation

The general protocol utilized to produce the Tag-Seq libraries is described in detail in (26). Total RNA was extracted with RNeasy Micro Kit (Qiagen) without the DNase treatment step and fragmented by hydrolysis. The polyA-tail bearing RNA species were captured with oligo-dT beads and ligated to P5 adapters containing moderately degenerate base regions (mDBR) to later allow demultiplexing and deduplication of the data. First-strand cDNA was obtained from an oligo-dT-VN primer and used as template in a 10-cycle PCR with one of the primers tailed with the second, regular index (see Table S1 for the list of the used P5 and P7 indexes). Libraries to be run on the same flowcell were pooled, co-purified and size selected in the range 270–500 bp on LabChip XTe with DNA 750 Assay Kit (PerkinElmer). The broad size range of the selected fragments was achieved by combining two narrower size intervals which resulted in two-peaked distributions of fragment sizes. The pooling was performed equimolarly, with the exception that for the embryonic stage the amount of the PCR product taken was increased 1.5 times relative to the other samples to increase sensitivity to the potentially subtler differences in expression levels.

The samples were processed in three batches, and in total three High-Output 75-cycles single-end runs were performed on an Illumina NextSeq 500 machine. Batch and run assignment for individual samples was randomized. See Supplementary File 1 for details on sample indexing and read numbers.

### Reference transcriptome

Reference genome assembly of the conspecific clone KAP4 of North American *D. pulex* (RefSeq assembly GCF_021134715.1) was used as the basis for gene quantification. To obtain a more comprehensive gene set, we recruited four unstranded RNA-Seq sequencing paired-end Illumina runs previously obtained by us from two bulk *Chaoborus* kairomone induction experiments on strain R9 (SRA numbers SRX1070836, SRX1070837, SRR32573570 and SRR32569538). The RNA-Seq reads were processed with fastp v. 0.23.2 (29) from trimming, deduplication and merging overlapping read pairs. The resulting read pairs, merged reads and singleton reads were mapped to the KAP4 genome with hisat2 v. 2.2.1 (30) and assembled with stringtie v. 2.2.3 (31) in the references-guided mode.

For the purpose of phylogenetic analyses, a clone-specific transcriptome assembly was obtained with spades v. 3.14.1 (32) using the fastp-processed RNA-Seq data from strain R9.

### Data processing and analysis

Cutadapt v. 1.9.1 (33) was used for quality, adapter and polyA trimming, retaining reads at least 35 nt long. Trimmed reads were demultiplexed and deduplicated with tagseq v. 0.2 (26).

Gene quantification was performed with Salmon v. 1.10.2 (34) using the transcript sequences from the reference KAP4 transcriptome extended by recruiting the R9 RNA-Seq data as described above. To define transcript groups distinguishable with the generated TagSeq data, we performed transcript clustering with Terminus v. 0.1 (35) with a minimum spread value of 0.05, tolerance of 0.01 and consensus threshold of 0.25. Transcript groups were named after the representative transcript with a suffix indicating the type of relationship within the group: “∼GN” for groups corresponding to entire genomic loci, “∼TR” for single transcript from loci in which different transcripts belong to different groups, “∼GNs” for groups covering all transcripts from more than one genomic locus and “∼TRs” for other, more complex cases.

Differential expression analysis was conducted with DESeq2 v. 1.42.0 (36) with likelihood ratio as the test statistic when assessing null hypotheses. To reveal the genes differentially expressed under both treatments (natural and synthetic kairomones), a two-stage testing procedure was implemented using stageR v. 1.24.0 (37). In the screening stage, likelihood-ratio test (LRT) was performed for each gene to test the full model (both kairomone treatments) against the intercept-only reduced model, done separately for each developmental stage. In the confirmation stage, LRT was performed to test the individual terms of the full model for the genes passing the screening stage with FDR cutoff of 0.05. For the analysis of treatments, the read counts were fitted to the negative binomial distribution independently for the four stages and rlog-transformed (36). Gene expression patterns were analyzed with DEGreport v. 1.38.4 (38).

### Gene function analysis

Two sources of gene function annotations were used: protein domains and families were predicted for KAP4 proteins with InterProScan v. 5.61-93.0 (39) and GO terms were obtained from UniProt (40). For the GO term assignment, all UniProt records belonging to genus *Daphnia* were searched with BlastP from NCBI blast v. 2.15.0 (41) and hits with E-value less than 1e-10, identity ≥90% and alignment coverage of ≥90% were retained to transfer the GO-terms from the UniProt records. Over-representation and gene-set enrichment analyses were performed with ClusterProfiler v. 4.8.1 (42).

### Reference branchiopod assemblies

For gene phylogenies and LTR prediction, representative genome and transcriptome assemblies from Branchiopoda were recruited. Whenever gene predictions were not available in the source, TransDecoder v. 5.5.0 (43) was used to predict coding sequences in transcriptome assemblies (including the R9 transcriptome assembly) and GeneMark-ES v. 4.62 (44) in self-training mode was used to annotate genes in genome assemblies.

### Phylogenetic analysis of insulin-like peptides

For phylogenetic reconstruction of *Daphnia*’s ILP-1/3/4 clade, amino acid sequences of *D. pulex* KAP4 ILP-1 (RefSeq accession XP_046446612.1), ILP-3 (XP_046448317.1) and ILP-4 (XP_046448221.1) were used as BlastP queries against the protein sequences from the reference branchiopod assemblies (E-value threshold of 1e-3). For each assembly, matching sequences at least 100 residues in length were extracted and clustered at 95% identity level to exclude minor allelic variants with CD-HIT v. 4.8.1 (45). The collected sequences were combined, aligned with MAFFT v. 7.520 (46) in automatic mode, trimmed with trimAl v. 1.4.1 (47) to exclude alignment position with ≥10% gaps and phylogeny was reconstructed with IQ-TREE v. 2.4.0 (48) with 1000 replicates for ultrafast bootstrap (49).

### Analysis of long-terminal repeat elements

To collect and annotate long-terminal repeat (LTR) elements related to the kairomone-responsive LTR element, the following strategy was deployed. The gag protein sequence of the kairomone-responsive LTR element from *D. pulex* strain R9 was used as a tBlastN query to locate scaffolds containing related *gag* genes in other Cladocera with an E-value threshold of 1e-8. gag matches passing the empirical bit-score threshold of 45 were used to extract the *gag*-containing regions by adding up to 3000 nt upstream and 12000 nt downstream to the coordinates of the tBlastN match and excluding too short fragments (<1600 nt). LTRs were annotated with LTRharvest (50) and both LTR elements and fragments lacking LTRs were annotated with LTRdigest (51), both from GenomeTools v. 1.2.1. For gene annotation with LTRdigest, protein profiles distributed via GyDB (52) were used, in addition to two lineage-specific protein profiles for gag and Asp protease, with a HMMER E-value threshold of 1e-5. To create the lineage-specific profiles, R9 gag and Asp protease protein sequences from the R9 kairomone-responsive element were used as queries in tBlastN searches against branchiopod assemblies, as described above for gag, and aligned parts of the subject sequences were extracted, aligned with MAFFT and protein profiles were created with hmmbuild from HMMER v. 3.4 (53).

For phylogenetic analysis, complete *gag* genes (with start and stop codons, without frameshifts and coding for proteins at least 150 amino acids) were extracted from the annotated branchiopod LTR elements. The resulting protein sequences were combined with outgroup sequences: Arc proteins from human, rat, mouse, chicken and fruit fly, and gag protein of the yeast Ty3 retrotransposon. The sequences were aligned with MAFFT and trimmed with trimAl to trim position with ≥10% gaps. In parallel, CD-HIT was used to cluster the sequences at 90% identity and the gag proteins coming from the most complete genetic elements per assembly were picked from the trimmed alignment as cluster representatives. Phylogeny was reconstructed with IQ-TREE with 1000 ultra-fast bootstrap replicates.

### Gene knock-down

Gene-specific double stranded RNAi probes (for the insulin-like peptide transcript and the suppressor of cytokine signalling transcript) were generated from *D. pulex* R9 cDNA using specific primers with T7 promoter overhangs (Suppl. Table 1). Potential off-targets were searched for using NCBI blast. dsRNA synthesis was performed using the T7 RiboMAX™ Express Large-Scale RNA Production System (Promega). The resulting probes, each exceeding 200 bp in length, were administered systemically with Lipofectamine 3000 transfection reagent (Thermofisher). As a control we used a 300 bp eGFP probe (Eupheria, Germany).

Two solutions were prepared according to the manufacturer’s protocol. Solution A consisted of 25.25 µL phosphate-buffered saline (PBS; pH 7.4, 0.1 M) and 0.75 µL Lipofectamine 3000 reagent. Solution B contained 16.6 µL dsRNA (1500 ng/µL), 8.4 µL nuclease-free water, and 1 µL Lipofectamine 3000 reagent. The two solutions were combined to form the transfection mixture.

The transfection mixture was added to designated wells of a 24-well plate containing 2 mL of Aachener Daphnia Medium (AdAM; (54)) or kairomone-enriched medium, which was prepared by exposing the medium to 10 phantom midge larvae fed with 100 *Daphnia* juveniles over 24 hours. We included three conditions: (1) a control without kairomones and without dsRNAi probes, (2) a kairomone treatment with the eGFP control dsRNAi probe, and (3) a treatment combining one of the target gene dsRNAi probes with kairomone stimulation. For each group, one *D. pulex* mother with embryos in the red-eye stage (28) was placed into a well. Once the neonates were released from the brood pouch, they were transferred individually into fresh wells containing the medium and the corresponding dsRNAi mixture. Neckteeth expression was observed in the first and second juvenile instars using a stereo microscope (Olympus SZX 16) equipped with a digital camera controlled by TSO Vidmess software (version 3). Neckteeth were scored according to the criteria described by Tollrian et al. (55).

Effectiveness of the knockdown was validated via reverse transcription quantitative PCR (RT-qPCR) of the genes of interest. For that, animals were exposed to the respective probes as described above and collected for qPCR in the first juvenile instar in RNA/DNA shield (Zymo Research). RNA was extracted using the Quick-RNA Microprep Kit (Zymo) according to the manufacturer’s protocol. qPCR reactions were run with a Luna reverse transcription qPCR kit (New England Biolabs, Germany) containing 10 ng/µl RNA. We performed three biological and two technical replicates using *Stx16* as a reference gene as described by Spanier et al. (56). ΔΔCT analysis was performed using the package PCR v. 1.2.2 which (57). Significant differences in gene expression were assessed using Student’s T-test.

## Results and Discussion

### Study design and overview of the data

A total of 72 Tag-Seq libraries were prepared with each sample corresponding to an individual *Daphnia* clutch assigned to one of the four developmental stages from late embryo to juvenile 2 and one of the three conditions: control and two kairomone treatments, natural and synthetic (Fig. 1a). After raw data quality filtering and demultiplexing, on average 13,130,897 ± 4,620,673 reads (±SD) were retained per sample. With the chosen filtering settings, 88.7 ± 3.6 % of the reads per sample remained after PCR duplicate removal, of which 93.7 ± 1.6% could be mapped to the reference transcriptome (Suppl. File 1, Suppl. Fig. 1). Analysis of the read counts assigned to spike-in transcripts, demonstrated that the approach was quantitative and reproducible (Suppl. Fig. 2).

### Sample clustering

Unsupervised clustering of the samples based on *Daphnia* gene expression levels showed that the prevailing factor explaining the between-sample variability was developmental stage (Fig. 1b). Each sample grouped within one of the four clusters corresponding strictly to the respective stage. Despite clustering together with the other e18 samples, sample e18fs1 appeared as an extreme outlier in principle component analysis (PCA) indicating that the corresponding clutch might have contained maldeveloping embryos and was excluded. Clustering analysis and PCA also revealed the jI42 stage to be heterogeneous with two subpopulations designated jI42.1 and jI42.2 (Fig. 1b, Suppl. Fig. 3a). While the first principal component in PCA corresponded to the divide between premolt (e18 and jI42.2) and postmolt (jI30 and jII56) stages, the jI42.1 samples occupied an intermediate position in this axis and their separation from the rest of the samples was further explained by the second principal component (see Suppl. Fig. 3a). Expression levels of two markers of the premolting phase: the chitin synthase 1 gene *krotzkopf verkehrt* (58) and the 7-dehydrocholesterol desaturase *neverland1* (59), demonstrated a sharp rise in expression levels in the premolt stages e18 and j42.2 stages contrasting with low background expression levels in jI30 and jII56, with the juveniles jI42.1 again occupying an intermediate position (Suppl. Fig. 3b). Taken together, this indicates that jI42.1 clutches represent a developmental stage less advanced than jI42.2, hinting at the possibility that at around this time point (78 hrs post-oviposition), *Daphnia* undergo a developmental transition leading to two distinct yet temporally very close subgroups.

No clear separation of the treatments within the developmental stages was evident in the clustering pattern of the samples, with the exception of the jII56 stage, which was subdivided into two groups: all synthetic kairomone-treated and one natural kairomone-treated clutches in one group and all of the control and the rest of the natural hormone-treated clutches in the other one, a division also explained by the third principle component in PCA (see Suppl. Fig. 3a). We also noticed that the jI42.1 and jI42.2 sub-stages included unequal proportions of treated and untreated clutches with only a single control jI42.1 clutch (see Fig. 1b and Suppl. Fig. 3a). To avoid spurious results, for the downstream analyses we only took the jI42.2 clutches to represent this stage. Overall, we conclude that kairomone induction does not lead to large perturbations in gene expression levels in the earlier stages. In the second juvenile stage, induction with the synthetic kairomone, and to a lesser degree with the natural kairomone, leads to a significant departure in the development of the juveniles from the control. Our results also hint at the possibility that the kairomone induction might delay the jI42.1>jI42.2 transition, although larger sample sizes are required to test this hypothesis explicitly.

### General patterns of kairomone-induced differential gene expression

Inclusion of two kairomone treatments, the natural kairomone and the synthetic kairomone, in the experiment allowed us to assess how similar the responses to the two treatments are and to isolate the effect of the kairomone induction on gene expression from influence of the other compounds produced by *Chaoborus*. Differential gene expression analysis was thus performed in a two-stage manner to maximize the detection of genes responding to both treatments by first testing the general effect of kairomone treatment and then testing which treatments were responsible for the effect (see Materials and Methods for details). This strategy yielded a total of 443 differentially expressed genes (DEGs) which responded to both kairomone treatments across the four stages (Fig. 2, Suppl. Fig. 4).

**Figure 2.**
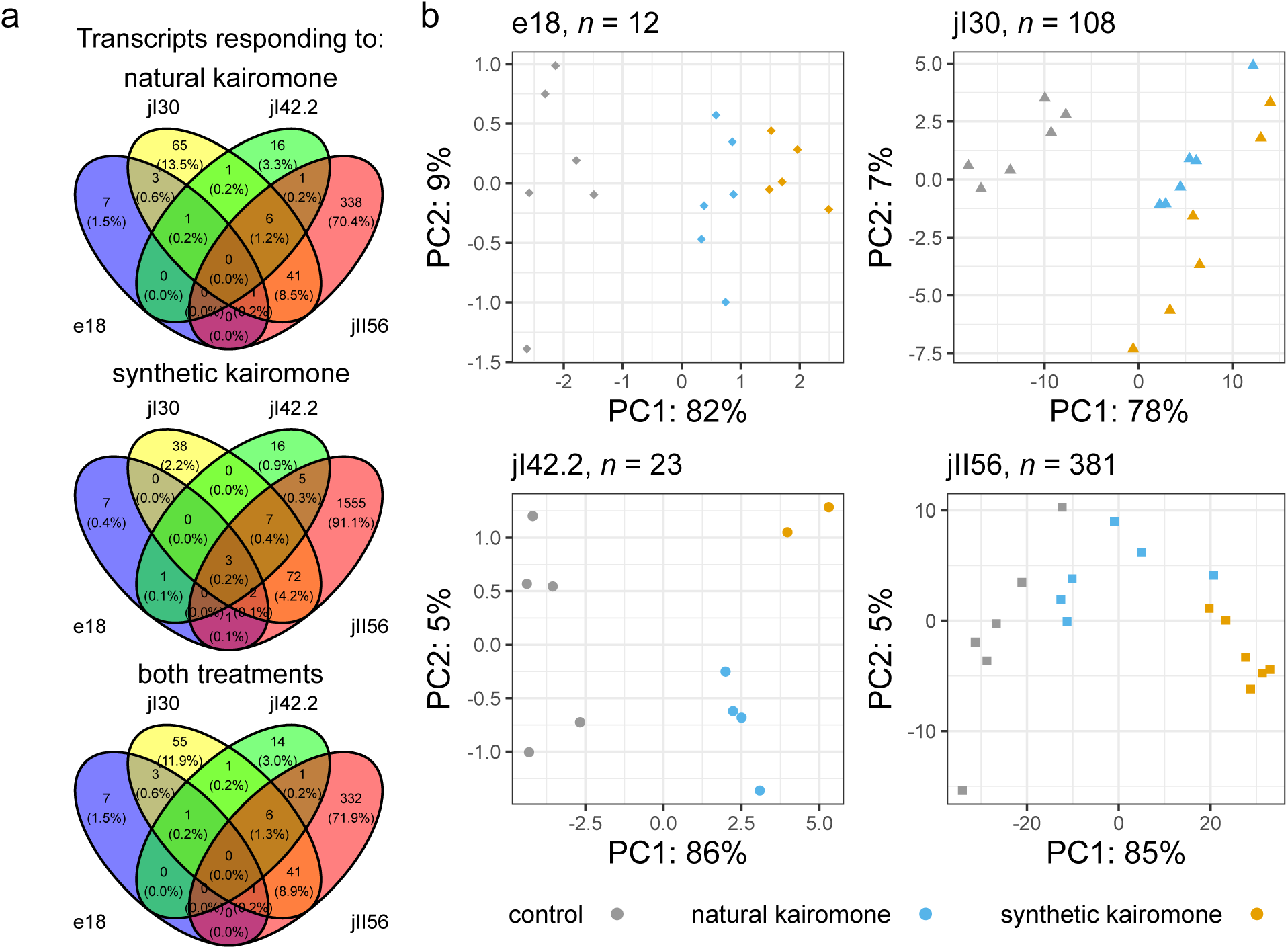
Summary of differentially expressed genes (DEGs) responding to the kairomone treatment. **a**, Venn diagrams showing DEGs shared between the four stages. Three partitions of DEGs are shown: those for which significant response could be detected in the natural kairomone or synthetic kairomone treatments and those found respond to both treatments. **b**, Principal component analysis based on the expression levels of DEGs for each stage, only the first two principal components are shown. Numbers indicate the corresponding numbers of genes per stage.

A considerable number of genes appeared differentially expressed only in one of the two kairomone treatments, with the synthetic kairomone demonstrating higher numbers, especially in the jII56 stage. Nevertheless, log-fold changes in the expression levels of the genes found to respond to both treatments were highly correlated between the two kairomone treatments in the same stage (Suppl. Fig. 5). The majority of the DEGs were upregulated and demonstrated a consistent direction of change across the different stages as indicated by high correlation coefficients in the log-fold changes of the DEGs shared by the corresponding stages (see Suppl. Fig. 5). Ordination of the samples based on expression levels of DEGs mostly demonstrated a clear separation of the samples into the three clouds corresponding to the three experimental conditions, yet the differentiation of the synthetic-kairomone-treated jII56 clutches was most distinct, mirroring the clustering pattern based on the overall gene expression levels (see Fig. 1a).

Noteworthy, the highest number of the differentially expressed genes was called in the latest stage, jII56, followed by the other post-molt stage, jI30. We hypothesize that there is a genuine tendency for the post-molt (jI30 and jII56) stages to have a stronger response to the kairomone treatments in comparison to the pre-molt stages (e18 and jI42). It is likely that for the late embryonic stage (e18), despite the higher number of read data generated, a disproportionate number of DEGs went undetected in our analysis due to small fold-changes and/or low expression levels, as at this early stage the kairomone response might be expected to be more subtle and restricted to particular organs or tissues. One compounding factor for the jI42 stage is the fact that only a subset of the samples was used for the differential gene expression analysis (see above), thus lowering the power to detect DEGs. Nevertheless, when comparing the apparent log-fold changes of DEGs between treatments and stages (see Suppl. Fig. 5), we noticed that the correlations between the fold-changes of genes found to be differentially expressed in stages other than jI42 and their fold-changes in jI42 were insignificant or weakly negative for both kairomone treatments. This indicates that most of the corresponding genes were indeed either non-responsive or weakly responsive to the kairomone treatments in jI42.

Several expression patterns of the DEGs across the developmental stage could be discerned with the four largest groups of genes demonstrating one of the molting cycle-dependent patterns in the treatment, as well as in the control clutches (Suppl. Fig. 6). At the same time, a minority of the DEGs showed clinal expression patterns across development (e.g. groups 11, 8, 4 and 14) or expression dynamics differing profoundly between the control and the kairomone treatments (e.g. groups 1 and 5).

### Differentially expressed genes called in multiple stages

#### Early-onset DEGs: insulin signalling and putative kairomone detoxification

Genes with differential expression at multiple stages are of particular interest as they might represent general regulators or indicators of stage-independent physiological responses to the kairomone (Fig. 3a). The most prominent of these were three genes with early onset of differential expression and gradual decline with age: (i) a gene for insulin-like peptide (*ilp-3*); (ii) an ortholog of suppressor of cytokine signaling 2 (*socs2*) and (iii) *ugt209B1* coding for a UDP-glucuronosyltransferase.

**Figure 3.**
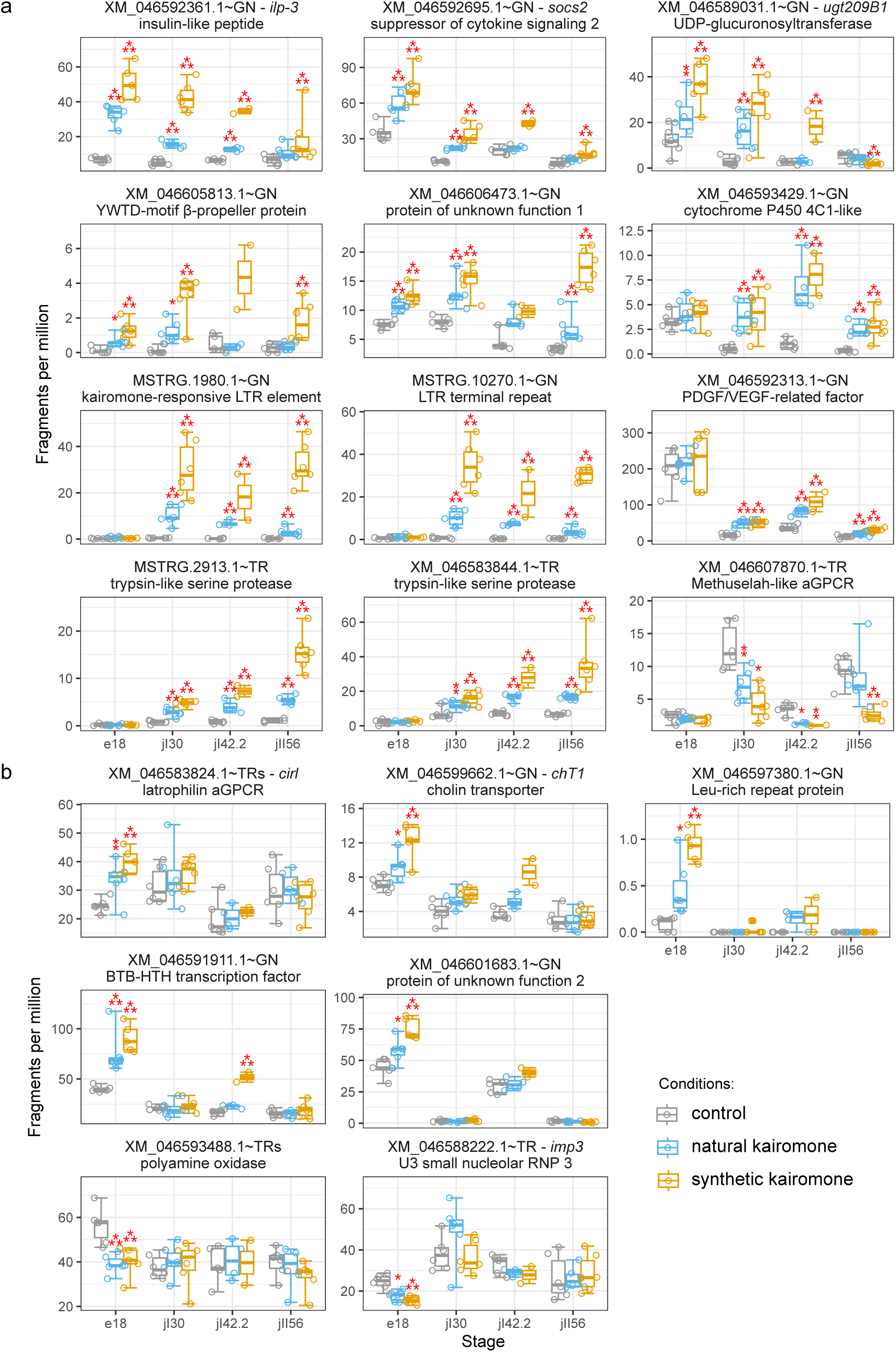
Expression levels of the top kairomone-responding genes. **a**, Transcripts found to be differentially expressed in at least three stages. **b,** Other transcripts differentially expressed specifically in the embryonic stage (e18). Asterisks mark level of confidence: ⁂ and ⁑ – likelihood ratio test (LRT) at the confirmation stage showing a significant contribution of the corresponding kairomone treatment to the full model with the overall FDR cutoff of 0.05 (⁂) or with raw *p*-value cutoff of 0.01 (⁑) and the shrunk log2-fold changes (LFC) exceed the threshold of 0.4; ⁎ – same as ⁑ but LFC lower than 0.4. All of the marked cases pass the screening stage with FDR cutoff of 0.05 and have at least one of the two LFC values (for the natural and/or synthetic kairomone treatments, respectively) exceeding the threshold of 0.4. Genes correspond to groups of transcripts which can be told apart using the Tag-Seq data (see Materials and Methods).

The first of these genes codes for the insulin-like peptide 3 (ILP-3 in (60), aIGF-2 in (61)) which has been implicated in the response to *Chaoborus* kairomone in our previous pilot RNA-Seq study (22). *ilp-3* was significantly up-regulated in both kairomone treatments during the first three stages: the expression differences were most prominent in the embryonic stage, but gradually decreased towards the final time point. ILP-3 belongs to the family of arthropod insulin-like growth factors (aIGFs) which in insects are produced by the fat body (61) and are known to be involved in phenotypic plasticity in insects, such as ornament development in the beetle *Gnatocerus cornutus* (62) and caste differentiation in the honey bee (63). We thus hypothesized that ILP-3 might be involved in the development of neckteeth as well. To test this, we knocked down *ilp-3* using double-stranded RNA interference (dsRNAi) (Fig. 4). The gene knockdown led to a significant decrease in the neckteeth expression with a particularly strong phenotype in the *ilp-3* knockdown juveniles of the 1st instar. The 2nd instar showed a much weaker effect of neckteeth reduction. Although a complete knockdown is not achieved with this approach (see Fig. 4c), these results suggest that ILP-3 might be involved in the control of neckteeth production, in particular in the first instar. Interestingly, *ilp-3* represents just one member of a small family of related genes in *Daphnia*. While the related species *Ceriodaphnia dubia* has one gene from this group, three IGF genes, *ilp-1*, *ilp-3* and *ilp-4*, are present in all *Daphnia* species resulting from two gene duplication events (Suppl. Fig. 7). High degree of synteny conservation between the genomic regions containing *ilp-3* in the neckteeth producing species, such as *D. pulex* and species producing other types of defensive structures or lacking them altogether, such as *D. magna* (see Suppl. Fig. 7), might suggest that this hormone is involved in systemic responses to kairomones across the genus and is not specific to neckteeth production.

**Figure 4.**
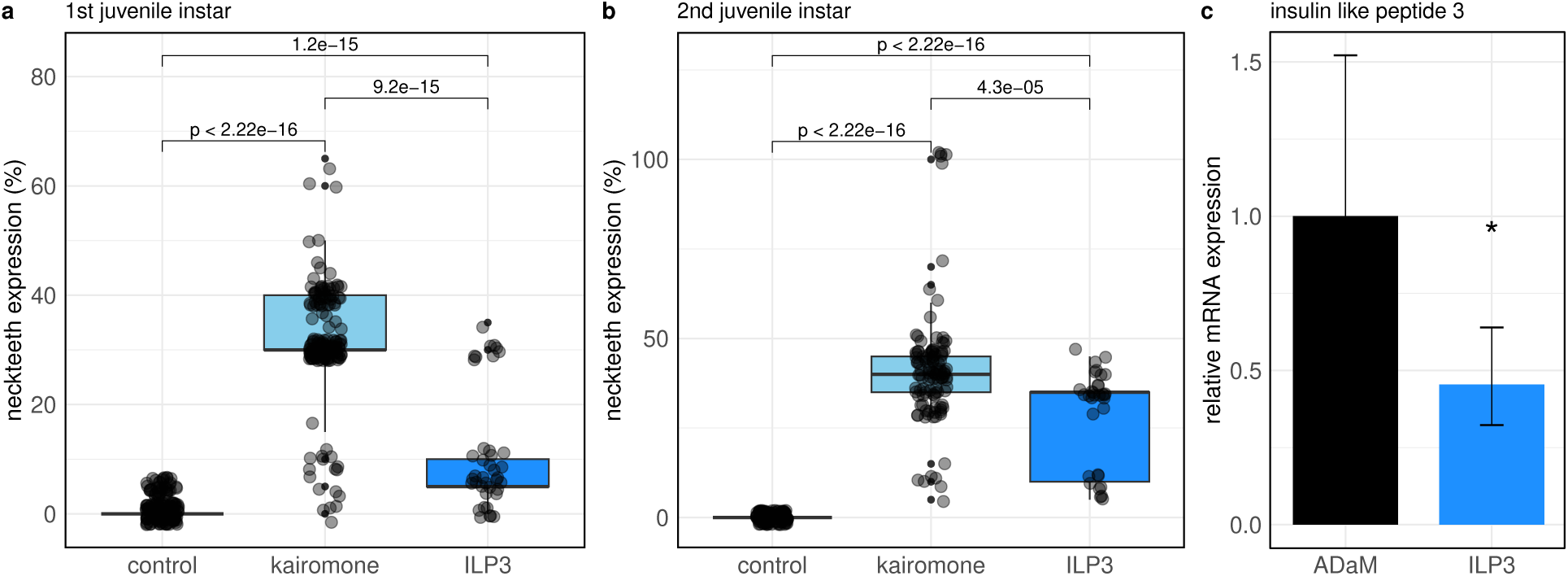
Knockdown of the kairomone-responsive genes *ilp-3* (insulin-like peptide) in *D. pulex* R9. **a,** Neckteeth phenotypes of the *ilp-3* in the first juvenile instar. **b,** Neckteeth phenotypes of the *ilp-3* in the second juvenile instar. Numbers indicate *p*-values of Bonferroni-corrected multiple comparisons using Dunn-Post Hoc Test. c, T-test results of qPCR checks for the expression levels of *ilp-3* in the control (ADaM) and knockdown juveniles. Asterisk indicates \**p*-values < 0.05.

We tentatively hypothesize that the second gene of this group, *socs2*, whose expression profile across stages and treatments closely matches that of *ilp-3*, might be functionally connected to it. In arthropods, SOCS2 has been implicated in a wide range of functions from innate immunity to modulation of ecdysteroid signaling and release of catecholamines (64–67), although the precise mechanisms of its action remain unknown. In vertebrates, SOCS2 has been demonstrated to be a negative feedback regulator of growth hormone/IGF-1 signaling in different tissues, facilitated by inhibition of STAT signaling and probably by binding to the IGF-1 receptor (68–73).

XM_046592313.1∼GN coding for another putative hormone, a PDGF/VEGF-related factor (PVF), might belong to the same network of morphogenetic factors, although its expression dynamics diverge from those of *ilp-3* and *socs2*: the sharp peak of its expression falls on the embryonic stage in both control and induced clutches (see Fig. 3a). This indicates that this PVF is also involved in morphogenetic processes directly unrelated to kairomone induction which in turn might have masked the more localized up-regulation of this gene in the kairomone treatments at this stage.

The third gene with an early onset of differential regulation, gradual decline and even down-regulation in juvenile 2 stage is *ugt209B1*. UDP-glucuronosyltransferase (UGTs) conjugate sugar moieties to diverse small lipophilic molecules and one of their typical functions is detoxification of xenobiotics (74,75). *ugt209B1* and several similar paralogs are related to the insect single-copy UGT50 family (76) although the multitude of these genes in *Daphnia* makes them more similar to the other multi-copy UGT families found in insects (77). Given its expression profile, one might expect UGT209B1 to be involved in kairomone detoxification. A connection can be made here with another gene likely involved in detoxification, XM_046593429.1∼GN coding for a cytochrome P450 4C1-like protein (see Fig. 3a). Nevertheless, given its dissimilar expression dynamics, later onset of differential expression and a nearly one order of magnitude lower expression levels, the oxidative reaction it performs might not be coupled with the conjugative reaction of UGT209B1.

Other genes with an early onset of relatively constant differential expression include two genes of unknown function: (i) XM_046605813.1∼GN coding for protein with a single YWTD-motif β-propeller domain and a single transmembrane helix but lacking other domains typical of lipoprotein receptors such as epidermal growth factor (EGF)-like domains (78) and (ii) XM_046606473.1∼GN coding for a protein with a conservative domain otherwise frequently associated with a C1q domain in related proteins from *Daphnia* (data not shown).

#### Late-onset DEGs: proteases, Methuselah and an LTR element

Several DEGs called at multiple stages demonstrate the opposite expression pattern of a later onset of differential expression and low expression levels in the embryonic stage, including such heterogeneous cases as: (i) two protease genes (see below); (ii) a systematically down-regulated transcript XM_046607870.1∼TR coding for a member of the Methuselah adhesion GPCR family similar to *Drosophila*’s *Mthl15* gene and (iii) two transcripts mapping to genomic regions related to long terminal repeat (LTR) elements.

Various members of the Methuselah family have been implicated in life history changes and stress resistance in insects (79,80) which leads us to the hypothesis that this gene is one of the regulators of life-history changes in *Daphnia* as well.

The case of the transcripts mapping to LTR elements is rather peculiar. Assembly and read mapping of RNA-Seq, demonstrated that the short transcript MSTRG.1980.1 is a fragment of a larger mRNA containing an entire LTR element with near-zero expression in the control and high expression levels in the kairomone-induced juveniles (Suppl. Fig. 8) and mapping of the Tag-Seq data was consistent with a polyA tail present at its 3’ end. This kairomone-responsive LTR element (KRLE) contains two open reading frames (ORFs) coding for: (i) a Ty3/Gypsy-type gag polyprotein with intact matrix, capsid and nucleocapsid regions and (ii) a retropepsin. The presence of the gag polyprotein and the lack of a pol polypeptide in this element indicate that it has the potential to produce virus-like particles (VLPs) in the cytoplasm but is not capable of autonomous retrotransposition. Phylogenetic analysis of gag sequences revealed a large clade of Ty3/Gypsy elements specific to Cladocera with varying gene architectures (Suppl. Fig. 9). While most of these elements show signs of gene degradation, the majority of them still maintain ORFs for components of the pol polyprotein. It is thus noteworthy that the closest relatives of KRLE from KAP4 and R9 are similar LTR elements without *pol* genes found in all other members of the *D. pulex* complex, although there are closely related more complete retroelements retaining ORFs for the reverse transcriptase and integrase in *D. pulex* PA42 and *D. galeata* as well. Expression of some LTR elements is known to be triggered by stress conditions in *Drosophila* in which they are hypothesized to play regulatory roles in tissue regeneration (81). Further, the lack of the *pol* ORF in KRLE is a characteristic shared with *arc* loci in dipterans and tetrapods. *arc*’s are Ty3/Gypsy-type *gag* genes domesticated independently in the two animal groups that form VLPs in neurons and regulate synaptic plasticity (82–84). The combined evidence thus points to the KRLE as an LTR element in the process of domestication involved in *D. pulex* in the kairomone response. The transcript MSTRG.10270.1 in KAP4 maps to a genomic region containing an isolated single copy of the terminal repeat closely related to that of KRLE.

### Kairomone induction in the late embryonic stage: neurohumoral regulation and patterning

Genes which appear to be differentially expressed in both of the kairomone treatments exclusively or preferentially in the e18 clutches (Fig. 3b) might be expected to have specific functions in the early response to the kairomone. Two such genes are constitutively expressed across the developmental stages, are up-regulated in the kairomone-treated embryos and are functionally connected to neurohumoral regulation: *cirl* coding for a latrophilin-family protein and *chT1* coding for a choline transporter. dCirl in *Drosophila* is expressed in the nervous system and is involved in modulation of mechanosensation and its silencing leads to hyperactivity (85,86). The involvement of latrophilins in behavioral regulation in both the fruit fly and vertebrates has been linked to dopamine signaling (86,87). Choline transporter, in turn, functions in choline uptake in the presynaptic terminal for acetylcholine biosynthesis. This echoes our previous physiological experiments (15) which demonstrated that dopamine plays a key role in linking sensory perception to adaptive morphological changes in *D. pulex*, while acetylcholine enhances sensitivity to environmental cues, particularly predator-induced defenses.

Two genes of unknown function: XM_046597380.1∼GN coding for a leucine-rich repeat (LRR)-only protein and XM_046601683.1∼GN, have an overall low expression level in j30 and jII56, but are over-expressed in the kairomone-induced e18 embryos and their expression levels are also elevated in the kairomone-treated jI42.2 clutches. Proteins with LRRs are known to have a wide range of physiological roles and among LRR-only proteins of known function, the product of XM_046597380.1∼GN shows a remote similarity to Oplophorus-luciferin 2-monooxygenase non-catalytic subunit and vertebrate carboxypeptidase N subunit 2 and is not directly related to any of the previously investigated groups of LRR-only proteins from decapods and other invertebrates implicated in innate immunity (88). It is likely that this protein is involved in an unknown enzymatic reaction as a non-catalytic subunit. XM_046601683.1∼GN codes for a protein from a *Daphnia*-specific family with no recognizable homology to other protein families.

A similar expression pattern was also observed for XM_046591911.1∼GN which codes for a protein with a broad-complex/tramtrack/bric-à-brac (BTB) domain and a divergent helix-turn-helix (HTH) DNA binding domain. Among *Dropsophila*’s BTB proteins, this domain composition is similar to that of the bab1 and bab2 developmental transcription regulators, although the HTH domain shows high similarity only to that of the lesser known BTB-HTH protein CG3726 expressed in embryonic brain and larval nervous system (89). XM_046591911.1∼GN is thus a prime candidate for regulation of the early patterning of the morphological defense.

Finally, two genes are down-regulated in the kairomone-treated e18 clutches: XM_046593488.1∼TRs coding for a polyamine oxidase (PAO) and *imp3* coding for the small nucleolar protein IMP3. PAOs influence the relative concentrations of polyamines which have been connected to a range of physiological processes in fruit fly development and shown to be spatially and temporally dynamic (90). Down-regulation of the PAO of unknown specificity in the kairomone-treated e18 clutches brings its expression to the level seen in the juvenile stages: this suggests that the concentrations of the corresponding polyamines normally differ in e18 from the later stages but exposure to the kairomone levels out these differences.

IMP3 coded by the second of the two genes is a conserved essential component of the eukaryotic rRNA maturation machinery (91). Interestingly, we find the *imp3* expression to be variable not only between the treatments but also across the developmental stages with its expression levels doubling after hatching. The lower levels of *imp3* in e18 would be expected to manifest in lower overall translation rates at this stage with respect to j30, and the kairomone treatment would thus exacerbate this effect. The transient down-regulation of *imp3* in the kairomone treatments is similar to the pattern of an immediate up- and later sharp down-regulation of its *D. melanogaster* homolog (CG4866) observed shortly after switching adult flies to a restricted diet: down-regulation of the many ribosome biogenesis genes was connected to the lifespan extension induced by this stress (92). It is nevertheless important to stress that we did not observe a similarly systemic down-regulation of ribosome biogenesis in the kairomone-induced R9 embryos and that *imp3* might have additional functions beyond rRNA maturation (93).

### Gene set enrichment and overrepresentation analyses: signatures of cuticle restructuring

For the analysis of higher-level patterns in functions of DEGs we performed enrichment tests using predicted protein domains and family assignments of the *D. pulex* proteins. Proteins with chitin-binding and protease(-like) domains and the corresponding protein families, such as insect cuticle protein and chymotrypsin family are among the most consistently appearing groups of protein in both, the gene set enrichment (GSE) and over-representation (OR) analyses (Fig. 5). The same functional groups appear to represent the most typical component of the proteome of *Daphnia magna* cuticle (94), thus indicating that these proteins in our kairomone induction experiment are related to structural modifications of the cuticle. Indeed, in *D. pulex*, besides the cuticular neckteeth, kairomone induction leads to such cuticular modifications as increased thickness and a higher number of layers in the procuticle, altogether leading to a higher cuticular stability (95). Similarly, the pillars connecting the two layers of integument of daphnia’s carapace are also increased in numbers and distribution density in kairomone-induced juveniles (96). Further indication of the functional connection between protease(-like) proteins and cuticle is the appearance of fusion proteins with a protease-like or protease inhibitor-like and a chitin-binding domains among differentially expressed genes, such as XM_046592105.1∼GN and XM_046584065.1∼GN (Suppl. Fig. 10). As can be seen from the results of the enrichment analyses, the protein signatures that can be linked to cuticle often demonstrate contrasting patterns of expression with some proteins having negative and some positive log-fold changes in response to the kairomone treatment in the corresponding stages. A closer examination of the expression profiles of these proteins indicates a strong dependence on the molting cycle with some proteins down- and some up-regulated at the stages at which the corresponding transcripts peak (Suppl. Fig. 10). We also noticed that proteins with Spaetzle-like domains found to be enriched in the GSE analysis in the e18 stage, and similarly proteins from the haem peroxidase family found in the OR analysis in the jII56 stage likewise also represent another two of the top five categories of cuticle-associated proteins in *D. magna* (94).

**Figure 5.**
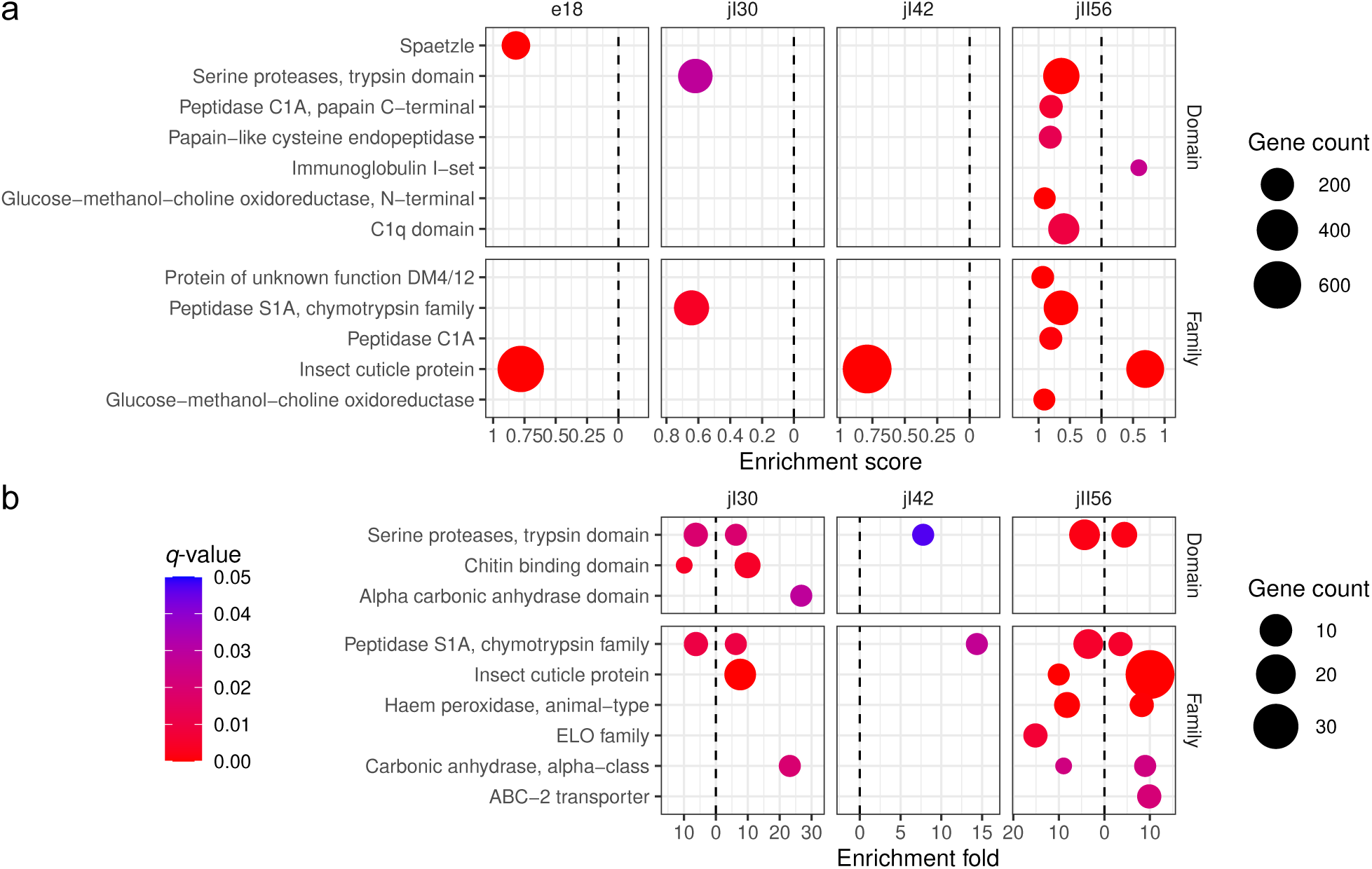
Enrichment analysis of protein domains and families among kairomone-responsive transcripts. **a**, Gene set enrichment analysis based on uncorrected log2-fold changes in the synthetic kairomone treatment. **b**, Over-representation analysis for genes responding to both kairomone treatments. The size of the dots reflects the number of distinct transcripts in the corresponding category. The color of the dots corresponds to the *q*-values. Dashed vertical lines separate down-regulated transcripts (left) and up-regulated transcripts (right).

These results indicate that the cuticle undergoes a profound restructuring and not a linear increase in abundance of its components. Moreover, this restructuring is stage-dependent as most of the genes in these categories appear to be differentially expressed at less than three stages and their expression levels follow the molting cycle (see Suppl. Fig. 10). The two related protease transcripts, MSTRG.2913.1 and XM_046583844.1, coding for proteins with trypsin-like domains represent an exception in this respect in showing a consistent increase in abundance in the kairomone treatments (see Fig. 3 and Suppl. Fig. 10) and are thus potentially not involved in the cuticle modification.

## Concluding remarks

In the current work we analyzed transcriptomic responses in *D. pulex* to its invertebrate predator, larvae of the phantom midge *Chaoborus*, using two experimental treatments: the *Chaoborus* kairomone as part of the natural mix of compounds emitted by the predator and the dominant component of the kairomone mix synthesized chemically (13). To our knowledge, this is the first study to analyze transcriptome responses of an aquatic organism to a kairomone synthesized *in vitro*. Our results demonstrate that the two kairomone treatments emit highly similar transcriptomic changes in *D. pulex* thus confirming the fact that fatty acid conjugates of L-glutamine excreted by the predator are responsible for triggering the complex anti-predatory responses in these crustaceans.

Gene expression analysis on the level of individual clutches across different stages, as opposed to large pools of individuals as used in previous experiments on anti-predatory transcriptome responses in *Daphnia* (18,20,22,23,97), provided us with the opportunity to study inter-clutch variation in gene expression across the different developmental stages. In this respect, the experimental design used here is very different from the one utilized in the non-replicated RNA-Seq study of An et al (23) in which control and *Chaoborus*-exposed pools of late embryos and juveniles spanning multiple stages were analyzed. On the other hand, analysis of individual clutches forced us to compromise between sample sizes and sequencing depth. This might be the reason behind the relatively low number of differentially expressed genes revealed in our study: the inter-clutch variation might have prevented us from assessing relatively subtle differences in gene expression levels. Our quantifications of differential gene expression should thus be considered conservative and to base our conclusions on a solid basis, we primarily focused on general trends in gene expression and on particular genes showing differential expression across multiple stages.

Our results demonstrate that gene expression patterns are highly dynamic in *Daphnia*’s development but that many of the genes involved in the transcriptomic responses to the kairomone production follow the molting cycle as well leading to a bias in the number of the differentially expressed called in the different points along the molting cycle. Consequently, we find an abundance of genes coding for components of the cuticle among the kairomone-responsive genes. The short list of the genes with constitutive differential up-regulation across the developmental stages include neurohumoral factors and detoxification proteins. Most interestingly, we find that elevated expression and gradual decline of the gene coding for insulin-like peptide ILP-3 is the strongest correlate of the neckteeth production in the juvenile stages thus bearing witness to the involvement of insulin-like signaling in the anti-predatory responses. Among the other neurohumoral factors implicated in neckteeth production in the previous physiological and candidate-gene studies (15,18,24,98), we find evidence for the involvement of neurotransmitter acetylcholine. There is little indication of the differential expression of enzymes involved in production of other neurotransmitters, such as dopamine or glutamate (15,23,24), or expression of their receptors in the earliest stages of kairomone response in our data. As indicated above, this might be connected to the relatively subtle changes in expression levels of these genes. Even if the exact connection remains to be shown, the up-regulation of the latrophilin gene *cirl* in the kairomone-induced embryos might be linked to dopamine signaling.

It is important to stress that the putative cellular determinants of neckteeth production are apparent in uninduced *Daphnia* (14), thus many components of the pathway leading to expression of this morphological hallmark of the anti-predatory defense, including the kairomone chemoreceptor, might be expressed constitutively and not change their expression levels upon exposure to the kairomone. Nevertheless, it is clear that our Tag-Seq approach has its limitations: increase in sequencing depth and number of samples will increase the list of detected differentially expressed genes, although to solve the more intricate tissue-specific changes in gene expression, single-cell transcriptomic profiling would be necessary. It is also important to stress that most of the *Daphnia* genes are functionally annotated based on homology to other model arthropods which together with the abundance of lineage-specific gene families and protein domains makes it difficult to draw conclusions about gene functions and pathways. We hope that the accumulating gene expression datasets, including the one presented here, will contribute to a better understanding of *Daphnia*’s gene functions and interactions.

## Data availability

The raw data were deposited in the NCBI SRA database (accession SRR4244255, BioProject PRJNA343077). Quantification results and differential gene expression results for analysis excluding jI42.1 clutches are included as Supplementary Files 1 and 2. Gene annotations, other quantification results and additional data are available via the Zenodo repository doi:10.5281/zenodo.15569083. The code used for the analyses is available at https://github.com/alephreish/kairomone-tagseq.

## Author contributions

AR, LCW and RT planned the experiment; AR performed kairomone-induction experiments for Tag-Seq and did library preparation; NK and UJ sequenced the libraries; TS and LCW did gene knockdown experiments and their analysis; AR performed the bioinformatic analyses and drafted the manuscript. All authors contributed to the discussion of the data and have read and approved the manuscript.

## Acknowledgments

The authors wish to thank Prof. Dr. Florian Leese (University of Duisburg-Essen) for kindly providing laboratory equipment for NGS library preparation. We thank Prof. Dr. Nils Metzler-Nolte (Ruhr University Bochum) for providing us with N-linolenoyl-L-glutamine. We are grateful to Isabel Schlurmann and Joshua Huster (Ruhr University Bochum) for performing preliminary gene knockdown experiments. The work was supported by the German Research Foundation (DFG) grant WE6019/2-2.

## Supplementary Figures

**Supplementary Figure 1.**
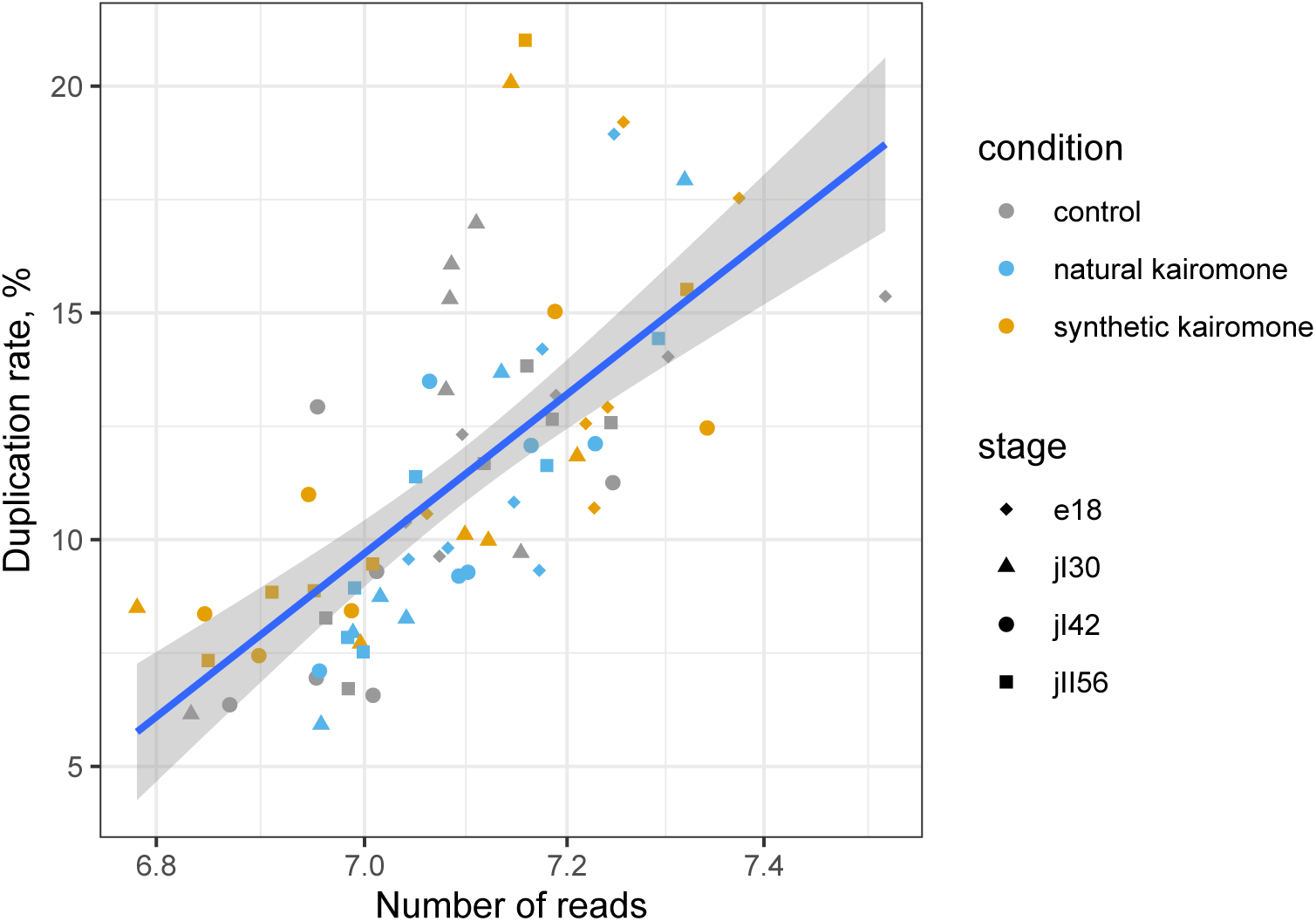
Proportion of PCR duplicates in the sequenced Tag-Seq libraries as a function of the number of the reads.

**Supplementary Figure 2.**
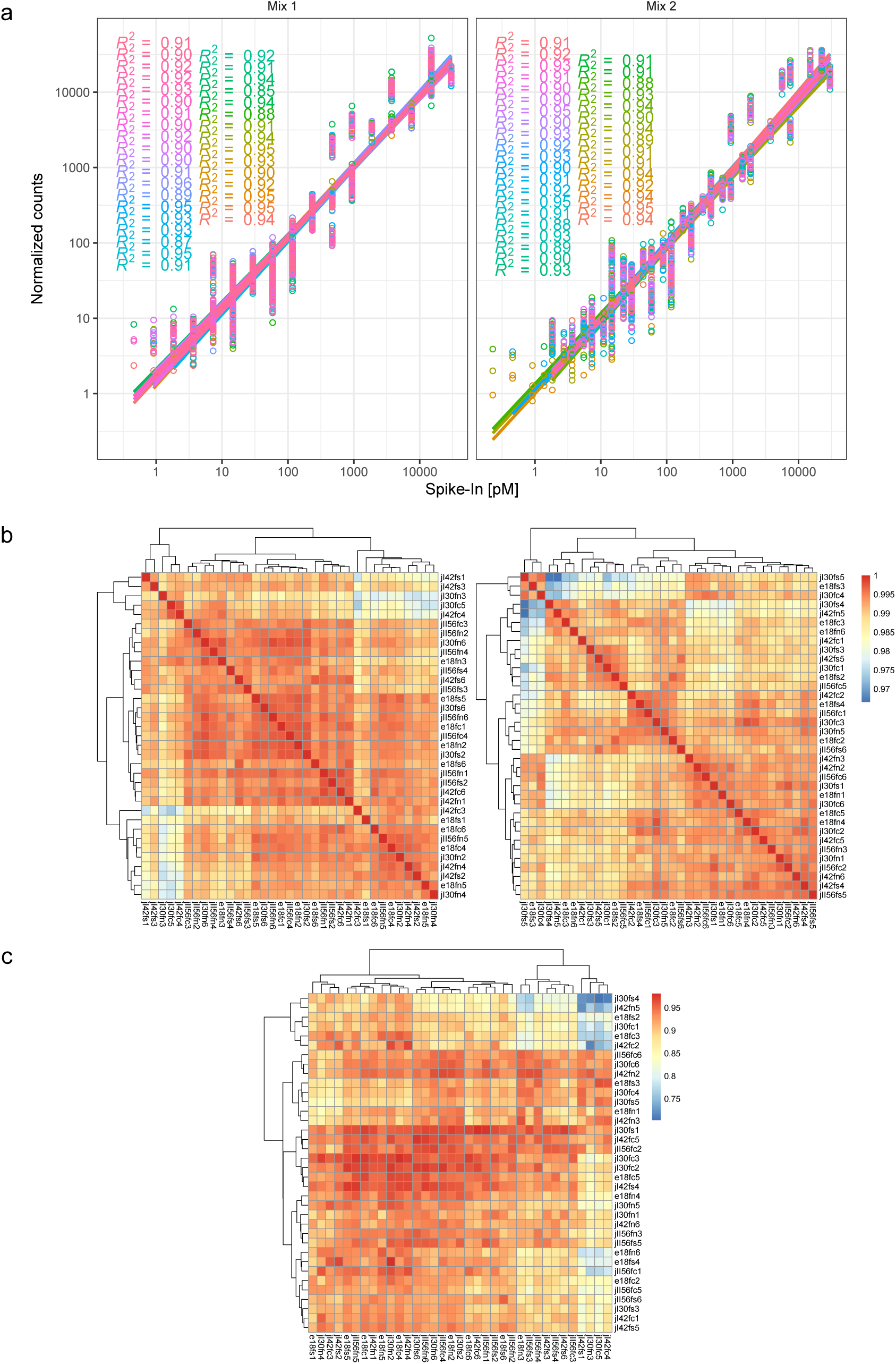
Quantification of ExFold spike-in transcripts in the Tag-Seq samples. **a**, ExFold transcript concentrations and their normalized Tag-Seq read counts. **b**, Correlations in the estimated ExFold transcript abundance between samples with the same mix (1 and 2, respectively). **c**, Correlations between log-fold differences in the abundance of ExFold transcripts predicted from pairs of samples assigned to mixes 1 and 2, respectively, and the expected log-fold differences. Only transcripts with at least 5 reads were considered.

**Supplementary Figure 3.**
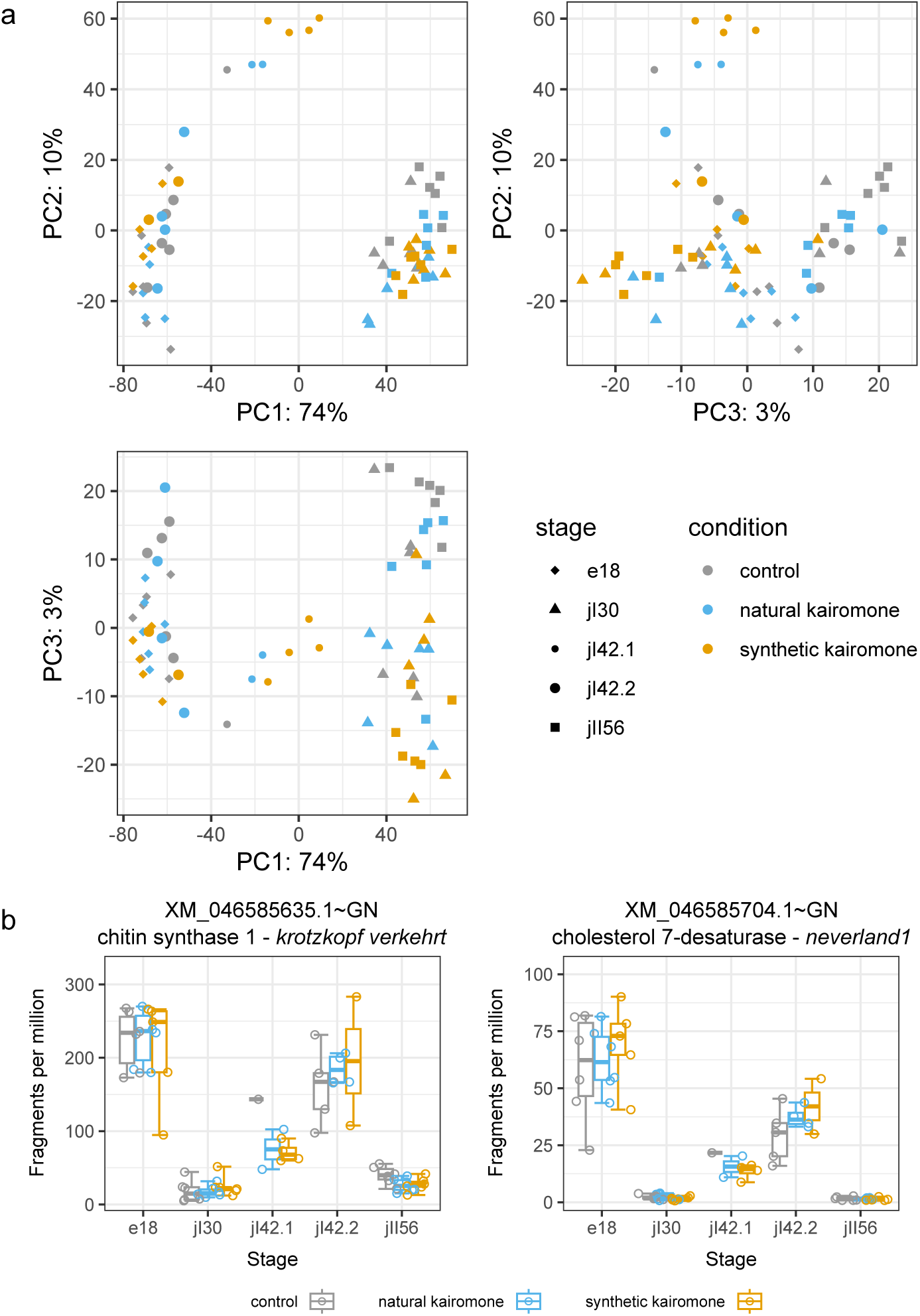
The main source of variation among the *Daphnia* clutches in the kairomone induction experiment is developmental stage and molting phase. a, Ordination of the Tag-Seq samples in the axes of principal components based on expression levels of the 500 most variable genes. b, Expression levels of the markers of molting cycle: chitin synthase 1 (*krotzkopf verkehrt*) and cholesterol 7-desaturase (*neverland1*). Whiskers denote min-max range, boxes — 25–75% quantiles, middle line — median.

**Supplementary Figure 4.**
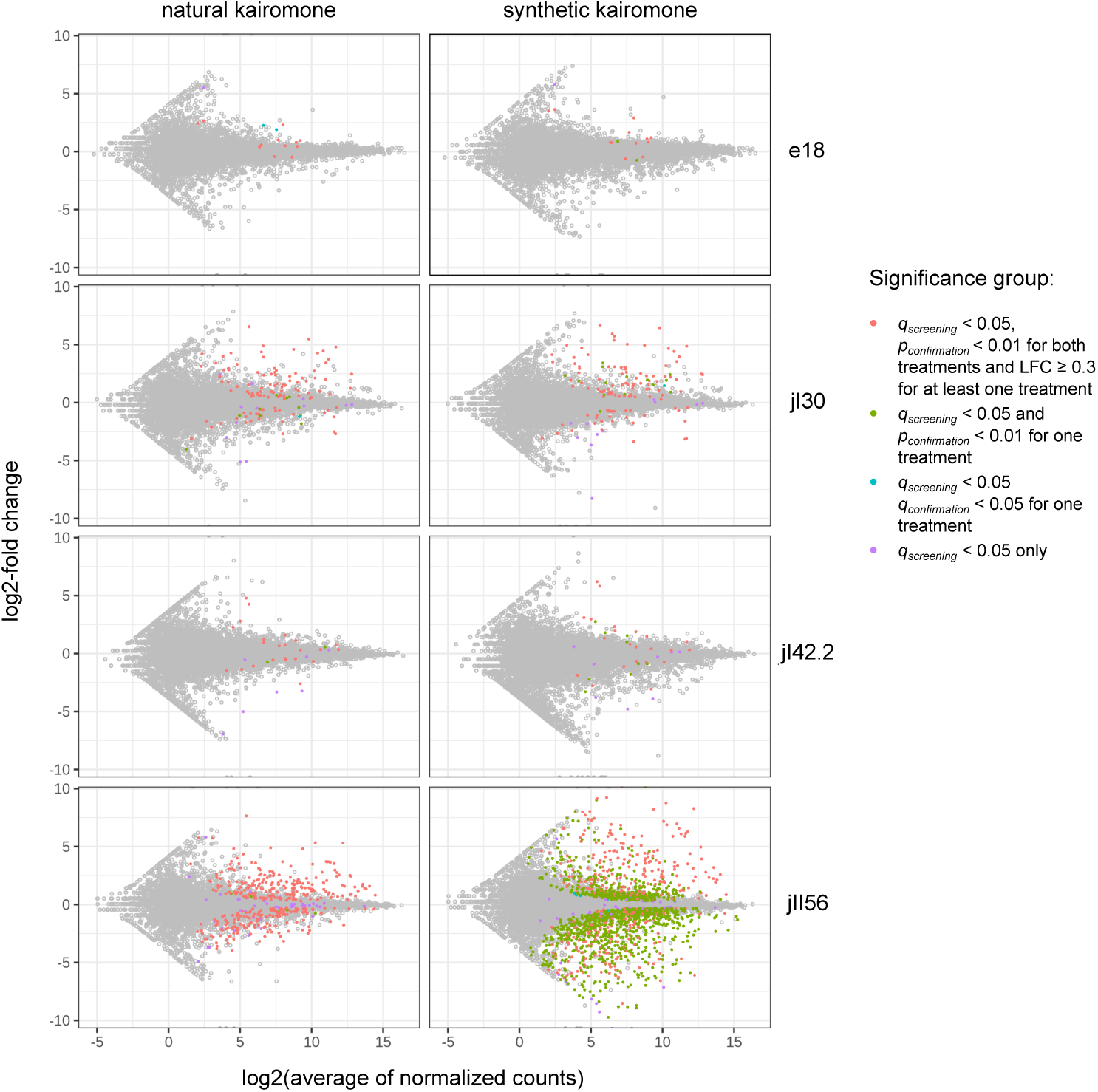
MA plots for two kairomone treatments across the four developmental stages. Each dot represents a gene (transcript group, see Materials and methods) with colored dots corresponding to genes passing one of the significance criteria as detailed to the right. *q_screening_*refers to the FDR-adjusted *p*-value in the screening phase, *p_confirmation_*and *q_confirmation_* refer to the unadjusted and FDR-adjusted *p*-values in the confirmation phase and LFC stands for log-fold change.

**Supplementary Figure 5.**
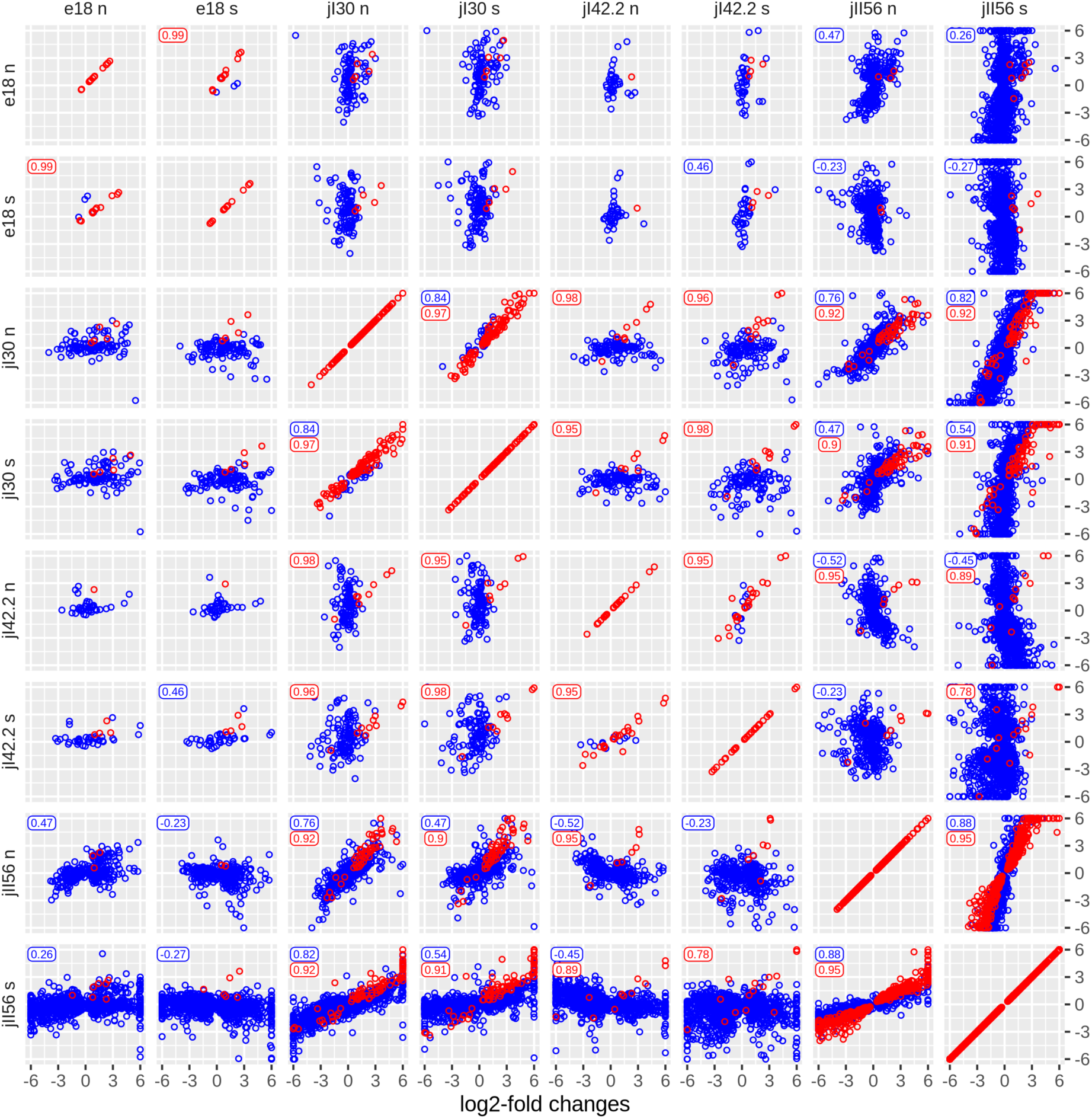
Correlations between unshrunk log2-fold changes (LFC) in expression levels of genes found to respond to the kairomone treatments. Each dot represents a transcript found to be differentially expressed in one (blue) or both (red) of the corresponding kairomone-stage combinations. For both groups of transcripts numbers in the left upper corner represent correlation coefficients when significant. LFC values exceeding 6.0 by module are capped.

**Supplementary Figure 6.**
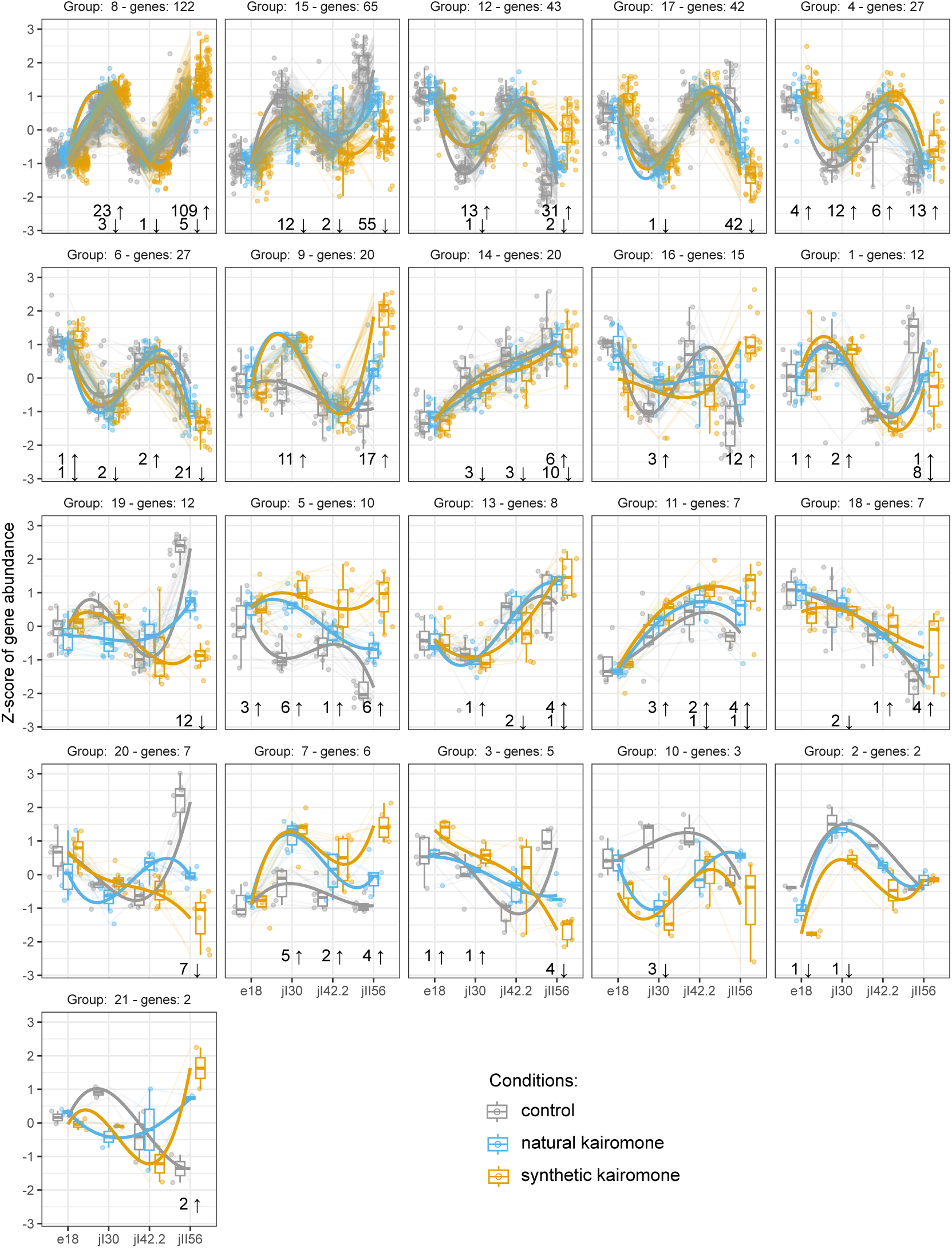
Expression patterns of the differentially expressed genes. rlog-normalized counts were used to group genes according to their expression levels across developmental stages and treatments. Numbers below the plots refer to the number of the genes differentially expressed in the corresponding stage and belonging to the group.

**Supplementary Figure 7.**
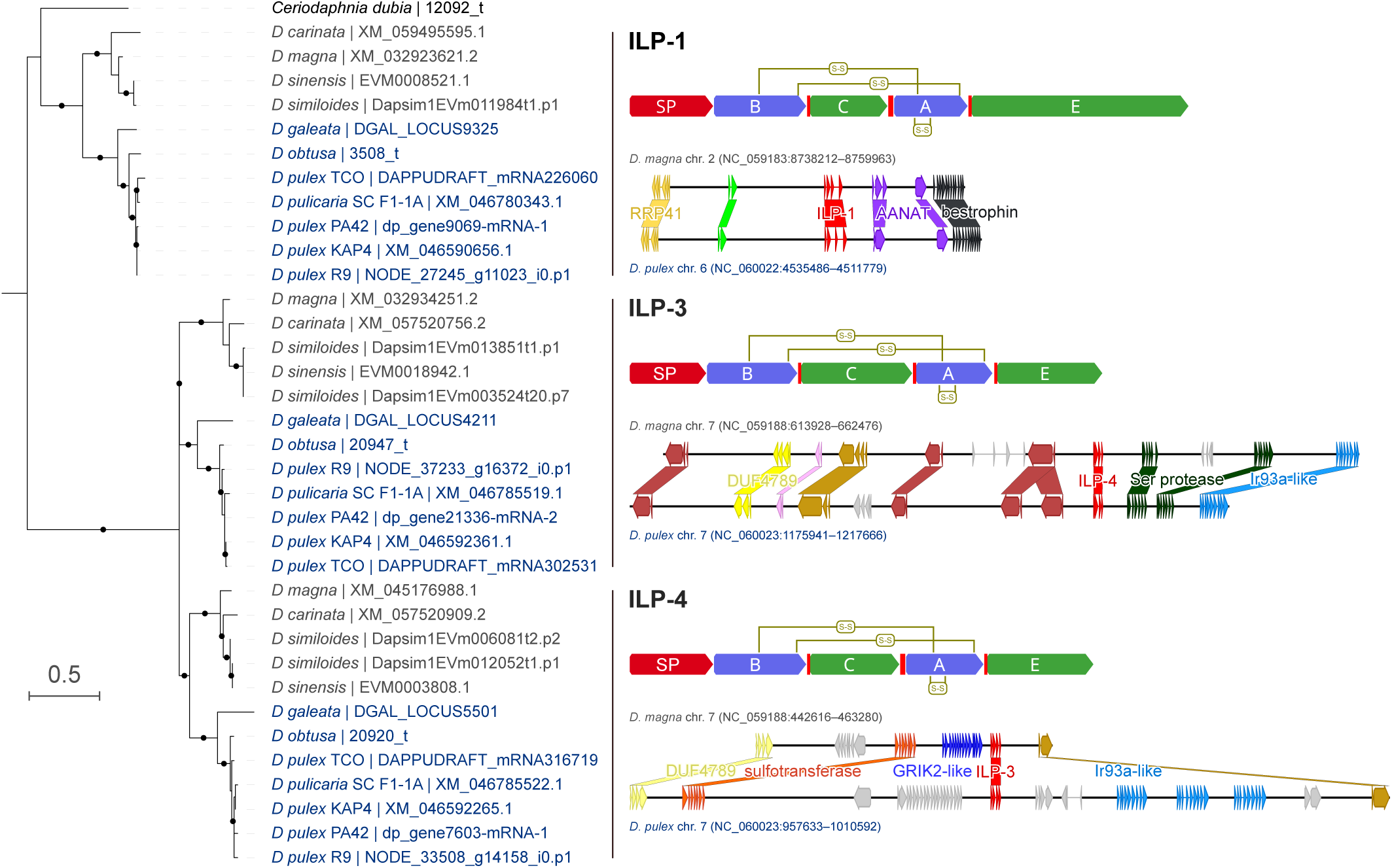
Arthropod insulin-like growth factors (aIGFs) in *Daphnia*. Left: Phylogenetic tree of the three aIGF genes, ILP-1, ILP-3 and ILP-4 in *Daphnia* (*Daphnia*) (blue labels) and *Daphnia* (*Ctenodaphnia*) (gray labels). Right: Predicted organization of the amino acid sequences of the three hormones and their genomic context with examples from the two *Daphnia* subgenera. SP — signal peptide, B and A — putative B- and A-chains, C — C-peptide, E — C-terminal extension. The intercalating red bars indicate potential cleavage sites at conserved mono- and di-basic residues. Predicted disulfide bridged between conserved cysteine residues are indicated as S-S (in yellow).

**Supplementary Figure 8.**
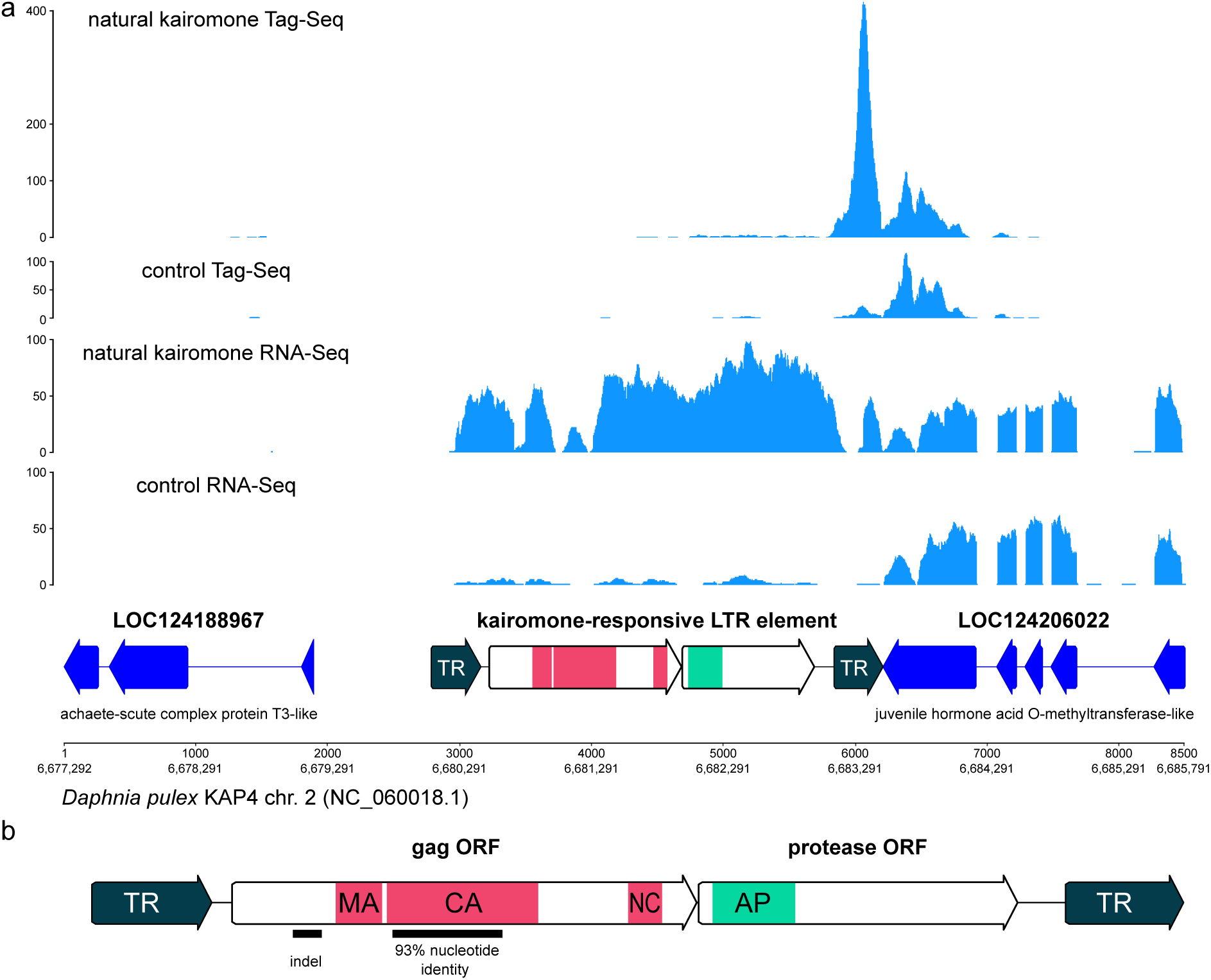
The *Chaoborus* kairomone-responsive LTR element. a. Genomic location of the LTR element (below) and the sequencing coverage around the locus in the Tag-Seq (exemplified here by merged data from jI30 control and natural-kairomone treated clutches) and RNA-Seq data (merged data from two pilot natural kairomone experiments with pooled juveniles 1). b. Structure of the LTR element with the terminal repeats (in orange) and the two ORFs. Homology regions are indicated for the ORFs: MA – matrix protein, CA – capsid protein, NC – putative nucleocapsid protein with Zinc-finger domains; PR – retropepsin-like aspartic protease. Indicated below are: an A-rich region which putatively enabled detection of transcription from the locus using the Tag-Seq protocol; two regions of lowered sequence identity between strains KAP4 and R9 responsible for the coverage gaps of the RNA-Seq data.

**Supplementary Figure 9.**
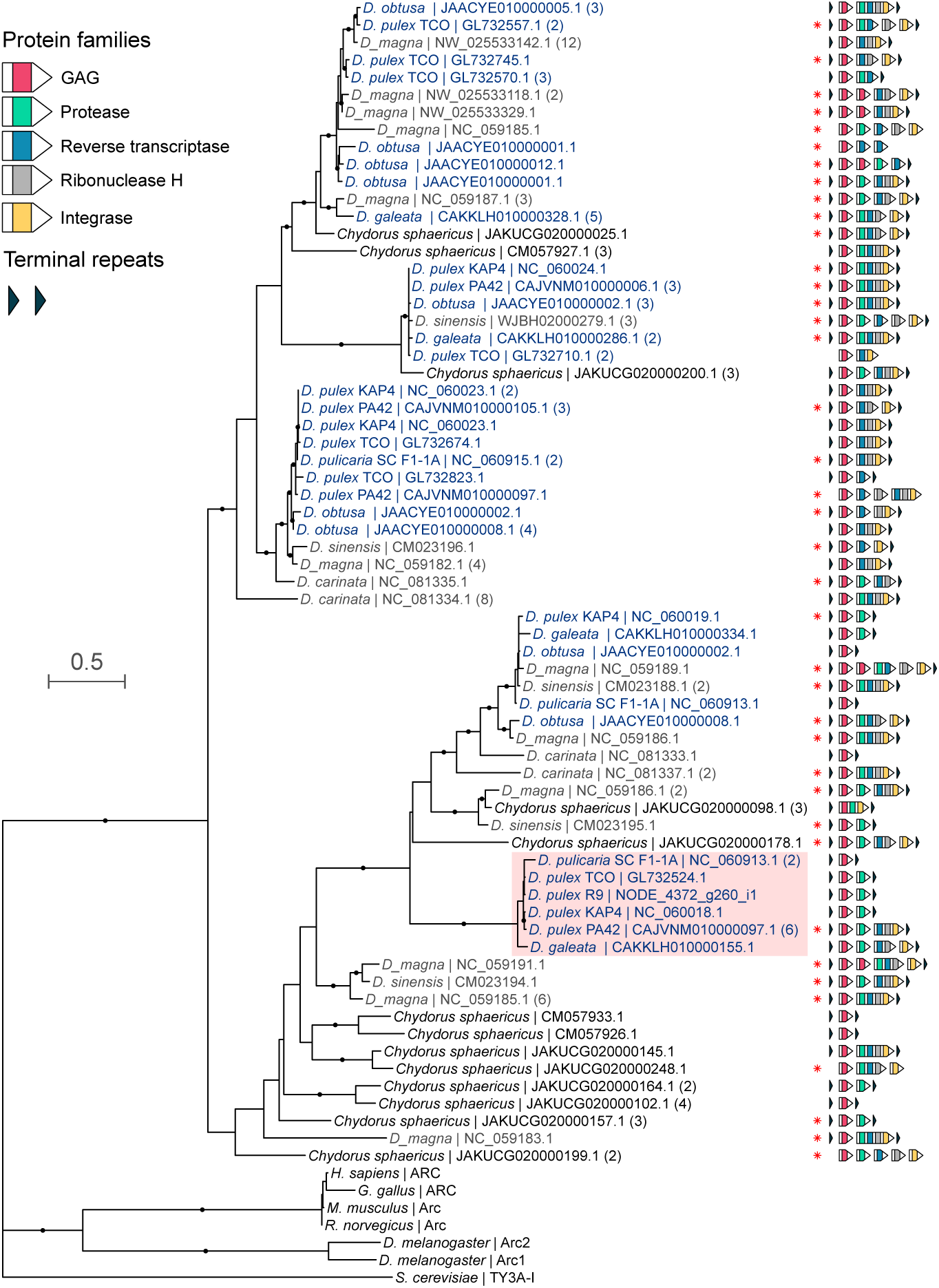
Phylogenetic analysis and architecture of the LTRs from a clade of retrotransposons related to the kairomone-responsive LTR element in *D. pulex*. Phylogeny is based on the representative protein sequences of the gag polyprotein. Representatives were chosen based on 90%-identity clusters (see Methods) and numbers in parentheses indicate numbers of cluster members. The color of the labels reflects subgenus: *Daphnia* (*Daphnia*) (blue labels), *Daphnia* (*Ctenodaphnia*) (gray labels) and other Cladocera (black). The structure of the corresponding loci is shown for each gag protein to the right: colored arrows indicate ORFs with the color showing the domain composition of the encoded proteins; black arrowheads indicate the presence of terminal repeats; red asterisks indicate elements with ORFs showing signs of degradation. The clade containing the kairomone-responsive LTR element in *D. pulex* R9 is highlighted in red. Branches with dots have ultra-fast bootstrap support values ≥90.

**Supplementary Figure 10.**
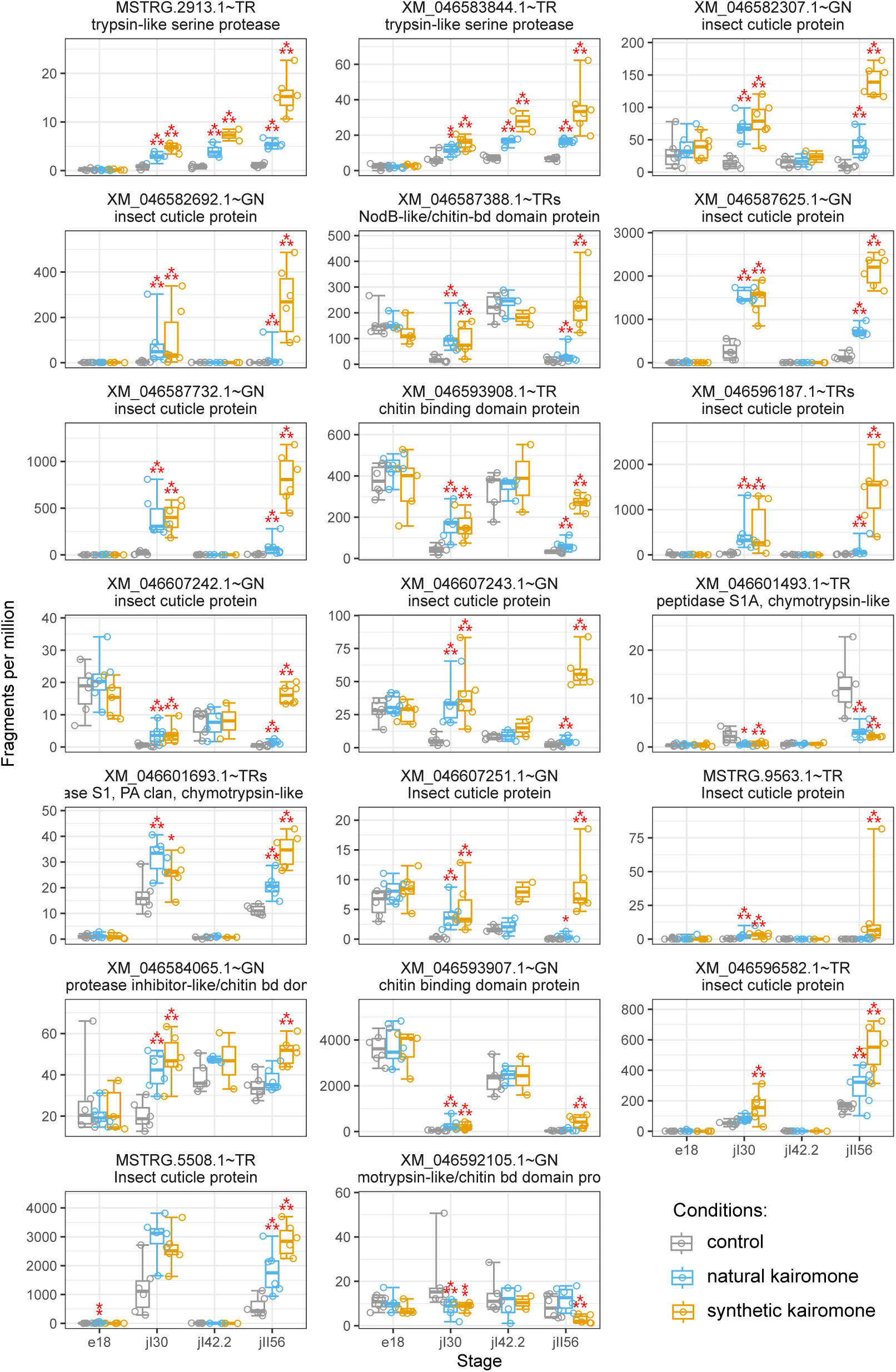
Expression profiles of genes coding for protease(-like) proteins, proteins with chitin-binding domain and insect cuticle proteins responding to the *Chaoborus* kairomone in at least two stages. Genes were selected based on the presence of InterPro signatures IPR001254, IPR000618, IPR043504, IPR001254, IPR001314, IPR002557 and IPR036508 and on significance of differential gene expression in both kairomone treatments in at least two developmental stages. See Figure 3 for explanation of the annotations.

**Supplementary Table 1.**
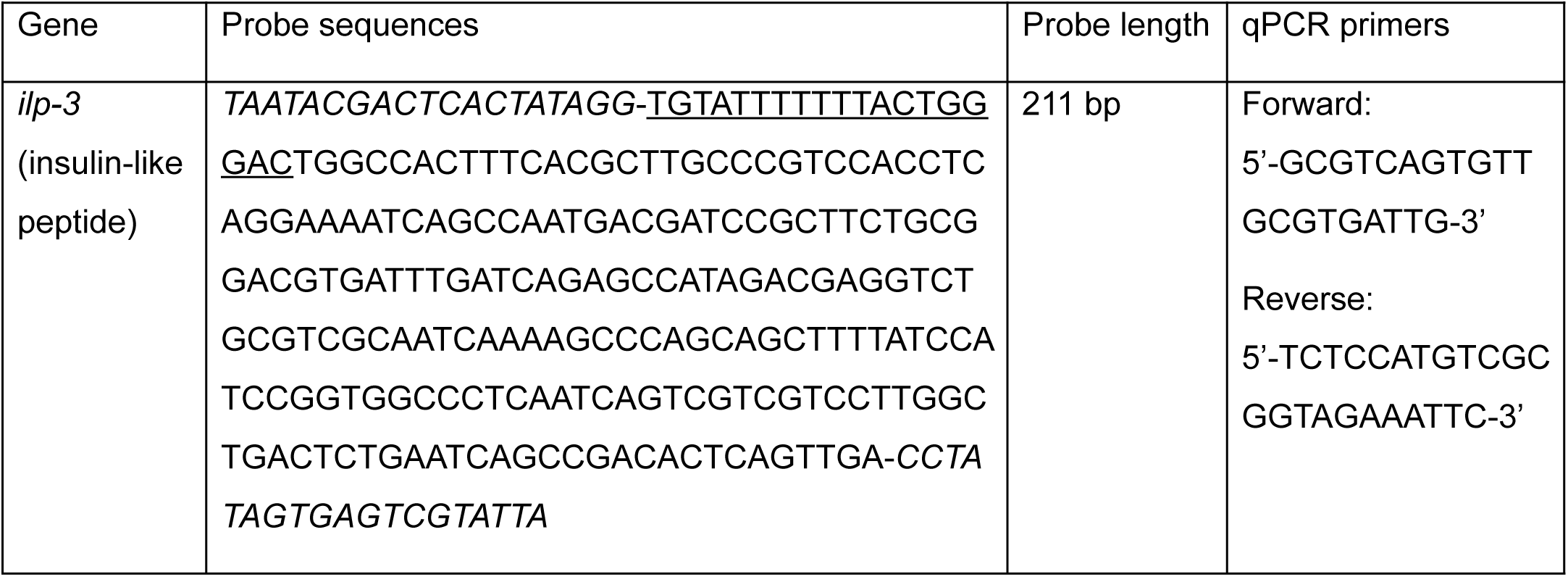
RNAi probes used for ilp-3 and socs2 knockdown and the corresponding qPCR primers used to confirm the knockdowns. Regions of the T7 overhangs are indicated in italics and primer regions are underlined.

